# Morphometrics of the preserved post-surgical hemisphere in pediatric drug-resistant epilepsy and implications for post-operative cognition

**DOI:** 10.1101/2023.09.24.559189

**Authors:** Michael C. Granovetter, Anne Margarette S. Maallo, Christina Patterson, Daniel Glen, Marlene Behrmann

## Abstract

Characterization of the structural integrity of cortex in adults who have undergone resection for epilepsy treatment has, in some cases, revealed persistent or even accelerated cortical atrophy but, in others, the converse is evident, and atrophy decelerates or even reverses. Whether this variability applies to a pediatric population, for whom postoperative plasticity may be greater than in adulthood, remains to be determined. Furthermore, understanding the morphometrics of this patient population is important, as cognitive gains have been associated with the anatomical status of preserved cortex post-resection. Here, we used high-resolution structural T1 magnetic resonance imaging data to compare the (1) gross anatomy, (2) cortical thickness, volume, and surface area for 34 cortical regions, and (3) volume for nine subcortical regions of 32 pediatric post-surgical cases and 51 healthy controls. Patients with either a preserved right hemisphere (RH) or left hemisphere (LH) had lower total white matter volume and select subcortical structures’ volumes, relative to controls; lateral ventricle size of both preserved RH and LH patients was also significantly larger than that of controls. However, relative to controls, only patients with a preserved RH had significantly lower total gray matter volume and lower thickness, volume, and surface area in multiple cortical regions, primarily in frontal and temporal cortex. The differences in preserved RH cortex of LH resection patients may relate to transfer of language function from the resected LH. Our findings lay the foundation for future studies probing associations of the morphometric differences in pediatric epilepsy surgery patients with neuropsychological outcomes.

## 1. Introduction

Epilepsy is a complex, progressive neurological disorder characterized by disruption of the normal balance of excitation and inhibition and sudden spikes of electrical discharge, all of which may result in neuronal impairment, axonal damage, and altered neural circuitry (Rossini et al., 2020). Pharmacologic treatment is often successful for seizure management, but, for ∼7-20% of pediatric and ∼30-40% of adult patients with drug-resistant epilepsy (DRE) (Tang et al., 2017; Xue-Ping et al., 2019), seizure control can be achieved by surgical resection of epileptogenic cortex. Post-surgical seizure freedom occurs in ∼86% and ∼81% of patients at 1 and 2 years follow-up, respectively (Weil et al., 2020) (see also (Jobst & Cascino, 2015)), and significant gains in several areas of cognition are reported in many individuals (Helmstaedter et al., 2020).

Because the post-surgical neuropsychological status relies largely on the post-surgical structural integrity of non-resected cortex, understanding the morphometrics of this residual tissue is important. Pre-surgical persistent and possibly accelerated reduction in cortical thickness (CxT) − a proxy for structural integrity of the brain (Federico & Wiebe, 2020) − has been identified beyond the defined epileptogenic zone/lobe and is evident even in the preserved hemisphere (Caciagli et al., 2017; Fonseca et al., 2022; Galovic et al., 2019; Grinenko et al., 2018), although this might be limited to a subset of regions. This reduction in gray matter (GM) appears to be ubiquitous across different epilepsy syndromes (Whelan et al., 2018). In contrast to these findings of ongoing structural compromise, post-surgical halting of progressive cortical atrophy has also been reported (Galovic et al., 2020; Galovic et al., 2019; Lopez et al., 2022; Park et al., 2022) and, dramatically, in some patients with temporal lobe epilepsy, the pre-surgical reduction in CxT can even be reversed to within normal limits (Galovic et al., 2019) (for commentary, see (McDonald, 2021)). Importantly, the extent of reversal is correlated with better seizure control (Federico & Wiebe, 2020; Galovic et al., 2020; Galovic et al., 2019), as well as positive cognitive outcomes (Skirrow et al., 2011). There is also reversal of not only reduced CxT, but also reduced cortical volume (CV), and this recovery, too, is positively associated with cognitive improvement post-resection (Zhao et al., 2021). Moreover, in an encompassing recent review of patients with temporal lobe epilepsy surgery, three possible interpretations of post-surgical changes are enumerated, including alterations that lead to damage and degeneration, recovery, or reorganization (Sainburg et al., 2025), but which of the three outcomes is to be expected and for which region remains to be determined.

One key limitation of understanding the morphometrics in the postoperative period is that most existing studies focus primarily on adults. A few pediatric studies have reported reduced CxT in the frontal lobe and in the right hemisphere (RH) in children with generalized epilepsy (Taherahmadi et al., 2025)) and diminished white matter structural integrity in temporal (and extra-temporal), especially in the fornix (Nguyen et al., 2011). An understanding of morphometric differences in children post-resection is of high priority. This is especially important given the greater potential for plasticity post-surgically in children than in adults (Fandakova & Hartley, 2020; Hubener & Bonhoeffer, 2014) and extrapolating from the findings from studies conducted with adults may be misleading. Studies of the anatomical status of post-surgical cortex are also sorely needed given the growing consensus that surgery is underutilized and should be considered a first-line intervention for DRE, especially in childhood (Consales et al., 2021; Cross et al., 2022; Vakharia et al., 2018).

Here, we compared the post-surgical morphometry of the preserved hemisphere in 32 children with DRE and 51 age-matched typically developing controls on (1) gross total volumes of the lateral ventricle (LV), GM, and white matter (WM); (2) CxT (the average thickness or distance between the white and pial surfaces), CV, and cortical surface area (CSA) separately for 34 automatically-segmented cortical parcels; and, last, (3) volumes of nine subcortical structures. CxT, CV, and CSA, in particular, are metrics that are commonly adopted in studying brain structure (Azzony et al., 2023) and have distinct genetic profiles, lifespan trajectories, links with neuropsychological factors, and disease associations (Frangou et al., 2021; Lyall et al., 2015; Wierenga et al., 2014). These metrics are also differentially associated with cognitive development and neurodevelopmental disorders (Fjell et al., 2015; Williams et al., 2023; Winkler et al., 2010): whereas CxT development − which has been associated with intelligence (Schnack et al., 2015) − follows a linear decrease with age (Sanders et al., 2022), CSA and CV follow a curvilinear trajectory with CV peaking earlier than CSA (Wierenga et al., 2014). The information that we can glean from these different measures thus offers unique and complementary insights. Moreover, although post-surgical reduction in CxT affects both hemispheres equally in adults (Galovic et al., 2020), here, we were interested in whether any post-surgical structural differences depended on the preserved hemisphere being left or right following childhood resection. Specifically, language acquisition and maturation is associated with increased cortical thinning in the triangular part of the inferior frontal gyrus (IFG) of the left hemisphere (LH) compared to the RH (Qi et al., 2019), and in those with language reorganization after left glioma, increased overall CV is noted in RH regions that are homotopic with LH language areas. Thus, the two hemispheres following childhood resection may be different than would otherwise be the case in adulthood once individuals fully acquire language (Pasquini et al., 2022).

Here, we focus our investigation and analyses solely on the preserved hemisphere, as any residual tissue in the hemisphere ipsilesional to surgery may have structural abnormalities either directly related to the surgery or because of ongoing epileptogenic activity. We employed general linear modeling (GLM) to compare gross anatomy, cortical morphometry, and subcortical morphometry of patients vs. controls and, separately, patients’ preserved LH vs. patients’ preserved RH. Because univariate analysis may be plagued by covariance between morphological variables (Wang et al., 2021), we also exploited multivariate analyses to identify models that include a combination of structural parameters and use these models to predict group assignment (patient vs. control) and, in the patients, any ongoing seizure burden, using the International League Against Epilepsy (ILAE) outcome classification schema (Fisher et al., 2017).

## 2. Materials and methods

### 2.1. Participants

We recruited 32 pediatric patients with DRE and subsequent cortical resection or ablation for this study. There were 13 patients with LH surgery and a preserved RH (median age/median absolute deviation of age: 15.7/1.7 years; 6 females, 7 males) and 19 patients with RH surgery and a preserved LH (median age/median absolute deviation of age: 15.4/3.7 yr; 11 females, 8 males). The patients’ demographic information and medical history are detailed in Table 1. Note that we use “RH patient” to refer to patients with left-sided resections but a preserved RH, as it is the RH from which we derive the measures, and the same holds true for “LH patient” in reference to the preserved LH of these patients.

**Table 1.**
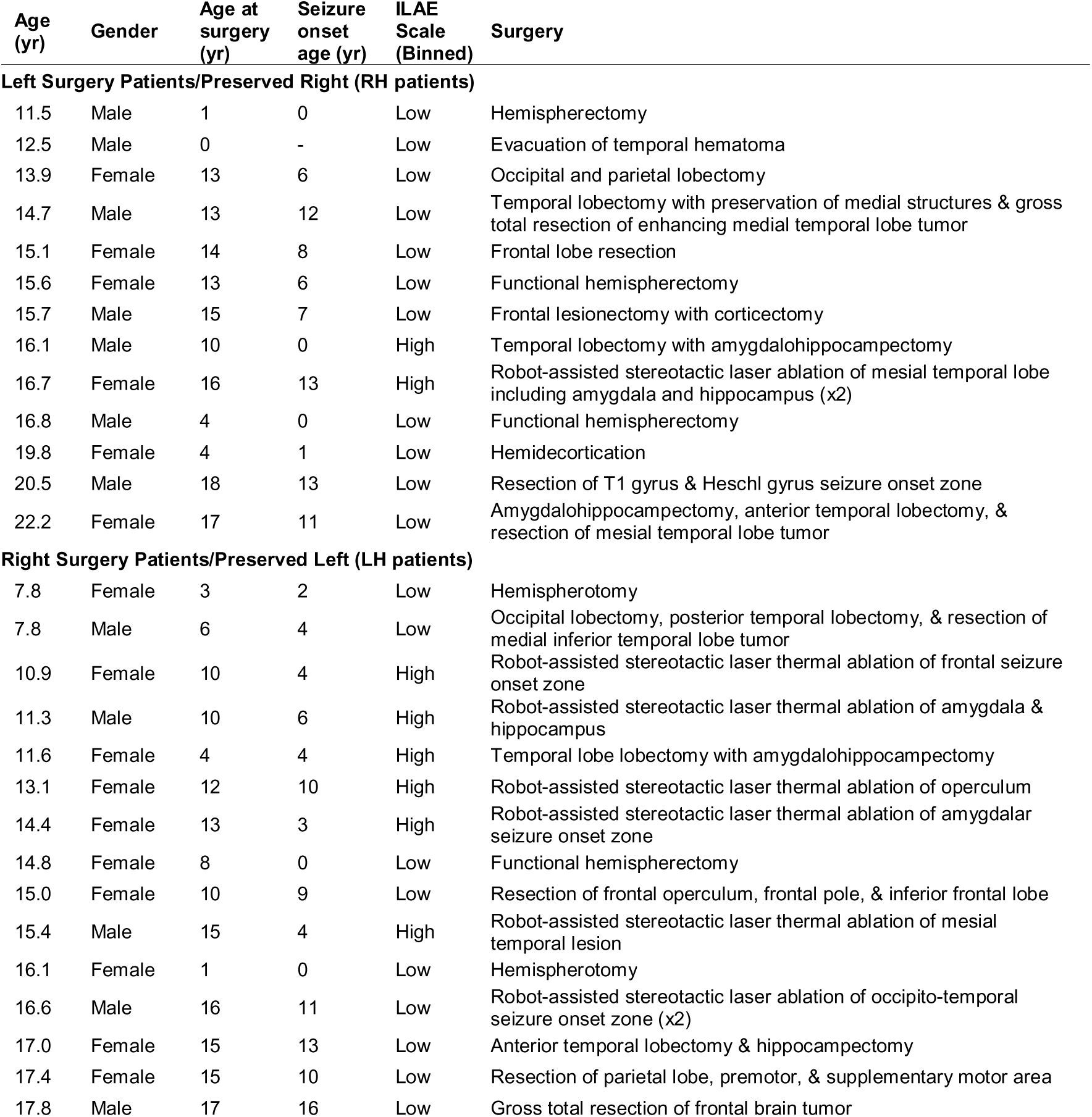
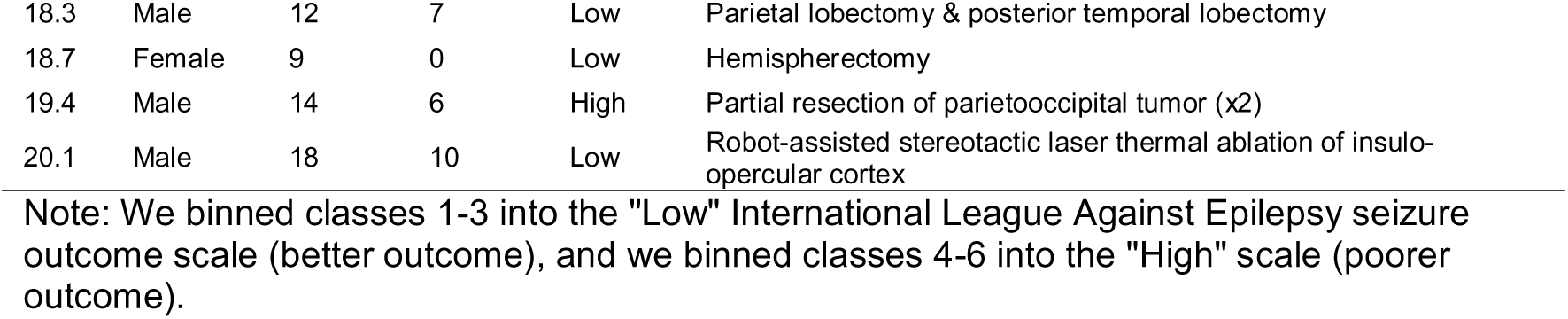
Patient information.

The two patient groups were matched on age at testing (*F*(1,30)=1.12; *p*=0.30), gender (*z*(30)=0.65; *p*=0.51), age at first surgery (*F*(1,30)=0.03; *p*=0.87), seizure-onset age (*F*(1,29)=0.01; *p*=0.93), and binned ILAE outcome scores (Scheffer et al., 2017) (*z*(30)=1.29; *p*=0.20). Most patients underwent surgery at the University of Pittsburgh Medical Center Children’s Hospital of Pittsburgh, and additional patients were recruited with the assistance of the Pediatric Epilepsy Surgery Alliance. Given the rarity of this patient population, we report retrospective analyses on patient scans from 2013 to 2022. We invited most patients for post-surgical scanning only.

The 51 control participants (median age/median deviation of age: 14.8/4.9 yr; 24 females, 27 males) did not differ from the patients in age, either in aggregate, (*F*(1,81)=1.18; *p*=0.28) or by hemisphere (LH: *F*(1,68)=0.17; *p*=0.69 | RH: *F*(1,62)=1.88; *p*=0.18), and we also matched controls to patients on gender (aggregate: *z*(81)=0.54; *p*=0.59 | LH: *z*(68)=0.80; *p*=0.42 | RH: *z*(62)=0.06; *p*=0.95). All participants received $75 per hour for their participation. Where possible, participants signed assent or consent, as appropriate, and guardians consented to a protocol approved by the Institutional Review Boards of Carnegie Mellon University (CMU) and the University of Pittsburgh.

### 2.2. MRI Protocol

We acquired T1-weighted images using an MPRAGE sequence (1 mm isotropic resolution, TE=1.97 ms, TR=2300 ms, total scan time≅5 min). We obtained images from either a Siemens Verio 3T scanner with a 32-channel head coil at CMU (36 controls, 13 patients) or a Siemens Prisma 3T scanner with a 64-channel head coil at the CMU-Pitt Brain Imaging Data Generation & Education Center (RRID:SCR_023356; 15 controls, 19 patients).

### 2.3. Outcome Measures

The anatomical images were preprocessed (motion correction, intensity normalization, and skull stripping) and then segmented with the recon-all pipeline of FreeSurfer (v7.1.0) (Reuter et al., 2010; Ségonne et al., 2004; Sled et al., 1998). We manually inspected the output. As the recon-all pipeline does not generally result in accurate segmentation of the brains of hemispheric surgery patients, for these large resection patients, the intact hemisphere was mirrored using affine and non-linear transformations (lesion_align program in AFNI) previously shown to be robust to anatomical aberrations (Glen et al., 2021; Maallo et al., 2020). This permitted the automatic segmentation of both the true as well as mirrored hemispheres, and only the data from the actual non-resected hemisphere were analyzed (for alternative approaches, see (Liu et al., 2021; Sainburg et al., 2025)).

Morphometric measures were derived from the preserved hemisphere of the patients and, separately, from each hemisphere of the controls, as follows: (1) gross measures of volume of GM, WM, and LV; (2) cortical morphometry for 34 Regions of Interest (ROI), parcellated according to the Desikan-Killiany cortical atlas (Desikan et al., 2006; Fischl, van der Kouwe, et al., 2004); and (3) subcortical structures (Fischl et al., 2002; Fischl, Salat, et al., 2004) of nine regions, with the volume of each structure normalized to the total hemisphere volume (e.g., percentage of volume of each structure relative to the sum of GM, WM, and LV volumes). For the cortical morphometry, for each participant, we calculated CxT in mm, which we did not normalize, as per standard guidelines (Westman et al., 2013); CSA in mm^2^ by taking the area of each region and dividing it by the mean of all of the regions in that hemisphere only; and CV in mm^3^ normalized by total hemisphere volume (Fischl & Dale, 2000; Han et al., 2006; Reuter et al., 2012).

### 2.4. Statistical Analysis

Data were analyzed in R (TeamCore, 2020) (for packages utilized, see Supplemental Table 1) and SPSS 29.0.1.0. To harmonize the data across the two MRI machines, ComBat (Fortin, 2021) was used to model the data as a linear combination of group, age, gender, and scanner, with the assumption that scanner effects have both additive and multiplicative factors (Fortin et al., 2018). Furthermore, for each measure and ROI, data were winsorized so as not to lose any data: we replaced values above the 95th and below the 5th percentile of the distribution with an approximation of the corresponding percentile values (separately for controls and patients, as well as by hemisphere).

Considering the relatively small sample sizes, for each dependent measure, we implemented permutation testing by randomly shuffling the group label, and we fit a GLM with group as the primary predictor of interest and age and gender as covariates. This was repeated 1,000 times to create a distribution of the β-coefficients for the effect of group. A *p*-value was then calculated as the percentage of occurrences in which the absolute value of the β-coefficient from the simulated distribution exceeded the absolute value of the true β-coefficient. Significance was ascertained at an α-criterion of 0.05, and, per measure, the Benjamini-Hochberg correction (Benjamini & Yekutieli, 2001) was applied to *p*-values across the total number of ROIs. In addition, for further adjudication of the findings, Bayes factor (BF) was computed by comparing the model with the group term to a null model without the group term (Lee & Wagenmakers, 2013; Wagenmakers, 2007). This was especially important to examine null results and to obtain further evidence to inform the absence of significant effects. BFs below 0.33 and above 3 were interpreted as evidence for the alternative hypothesis and for the null hypothesis, respectively.

Thereafter, to elucidate which measures, alone or in combination, predicted group membership for patients vs. controls and, thereafter, just between patient groups, we conducted forward binary logistic regression analyses. Such a procedure yields those variables that account for significant variance, over and above any significant variable/s already included in the model. Last, we predicted ILAE outcome score (high vs. low) for the patients using a similar multivariate approach. The *r^2^* value of the regression models are provided using the well-established Nagelkerke measure (a modification of the Cox and Snell R Square, which is derived from the likelihood ratio test statistic (Nagelkerke, 1991)). The *r^2^* value reflects the normalized statistic (scaled between 0 and 1) for the goodness-of-fit of the logistic model, making interpretation of outcome easier than typical regression analyses. Instead of obtaining an absolute *p*-value, the Nagelkerke procedure indicates that a value below 0.2 indicates a weak relationship between the predictors and the outcome, a value of 0.2 to 0.4 indicates a moderate relationship, and a value of 0.4 or higher indicates a strong relationship. Because we are only interested in values indicating at least a moderate or a strong relationship, we only report those computed *r^2^* values greater than 0.25, and, for those below 0.25, we simply report the presence of a weak relationship between the predictors and outcome and do not offer any further interpretation.

## 3. Results

For all analyses, we report the statistical comparisons in the following sequence. First, we report the findings of each GLM, separately for each of the patient groups relative to the hemisphere of their corresponding matched controls. We then describe the findings from comparisons of the patient groups against each other. Note that age and gender are included as covariates in every analysis. Thereafter, for each comparison, we report the results of the binary logistic regression analyses identifying the factors that best predict each group and ILAE assignment.

### 3.1. Gross Anatomical Morphometrics

GLM revealed that, relative to the LH of the controls, LH patients had significantly larger LV volume (*p*=0.02, *BF*=0.57), significantly smaller WM volume (*p*=0.05, *BF*=1.02), and no difference on GM volume (*p*=0.27, *BF*=4.37). Relative to the RH of the controls, the RH patient group, like the LH patients, also had significantly larger LV volume (*p*<0.01, *BF*=0.01) and significantly smaller WM volume (*p*<0.01, *BF*=5.56*10^-2^), but they also had significantly smaller GM volume (*p*<0.001, *BF*=1.6*10^-3^). Across the two patient groups, neither the volume of the LV (*p*=0.55, *BF*=4.54) nor the volume of the WM differed (*p*=0.13, *BF*=1.39). However, the LH patient group had significantly greater GM volume than the RH patient group (*p*<0.01, *BF*=1.98*10^-2^). See Figure 1 for these findings.

**Figure 1:**
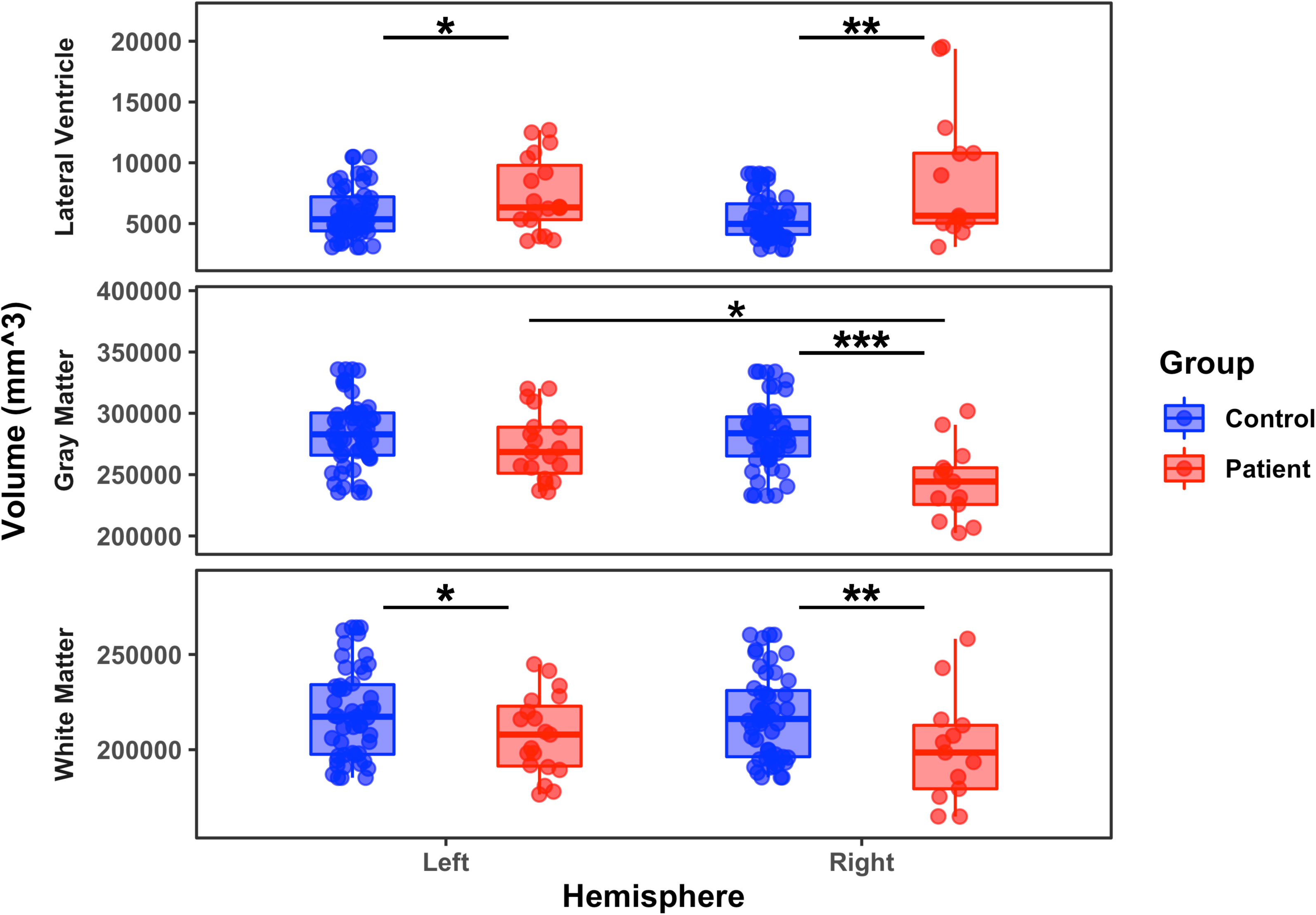
Volume of (top) lateral ventricles, (middle) gray matter, and (bottom) white matter for patients and controls, separately for the preserved left and right hemispheres. *: *p*<0.05; **: *p*<0.01; ***: *p*<0.001. Each dot represents a single participant, and the solid horizontal line in the box indicates the median of the group.

To evaluate which of these three dependent measures, LV, GM, and WM, singly or in combination predicted patient/control group membership, we used forward binary logistic regression with *p*<0.05 for entry criterion. LH patients and LH controls could not be reliably discriminated (model with low *r^2^* indicating a weak relationship between the predictors and outcome). This same analysis with data from RH patients and RH controls yielded a model that included GM and LV, in that order, but not WM, with an *r^2^* of 0.5 (strong relationship). In the direct comparison of the patient groups, GM was the best predictor of preserved hemisphere with *r^2^*=0.28 (moderate relationship). Last, we examined whether these gross anatomical measures predicted patients’ seizure burden post-surgery and found that no predictor was able to differentiate the ILAE binned outcome scores of the patients.

Patients with a preserved LH differed from their matched controls on LV and WM, but not GM (although these group differences are not completely well-supported, as we do not replicate them on binary regression analysis). The patients with a preserved RH differed significantly from controls on all three measures of gross anatomy; however, just GM and LV, and not WM, combined into a strongly predictive regression model. Across the two patient groups, GM alone moderately predicted patient group membership, and ILAE outcome could not be predicted by any dependent measure/s.

### 3.2. Cortical Regional Morphometrics

This section presents the GLM analyses separately for three dependent measures: CxT in mm, SA in mm^2^, and CV in mm^3^ for each of the 34 parcellated ROIs. We then report forward binary regression analyses, as above, separately for the three dependent measures, to determine the model/s that best predict group membership and the binned ILAE outcome scale. Last, we conduct a full multivariate regression in which we include all three dependent variables for each of the 34 regions in the models predicting group membership for each of the LH and RH patient groups against their respective controls, and then, predicting membership between the two patient groups.

#### 3.2.1. Cortical Thickness

As shown in Figure 2, relative to the LH of controls, the LH patients showed no statistically significant differences in CxT in any of 34 areas. In stark contrast, compared to the RH of controls, the RH patients had significantly lower CxT in 15 of the 34 cortical regions, and there was no region with significantly greater CxT for the patients than controls. Between the patient groups, the RH patients had lower CxT than the LH patients, but this held statistically in only three regions: the caudal middle frontal, rostral middle frontal, and superior frontal regions (see Supplementary Table 2 for individual *p* and *BF* values).

**Figure 2:**
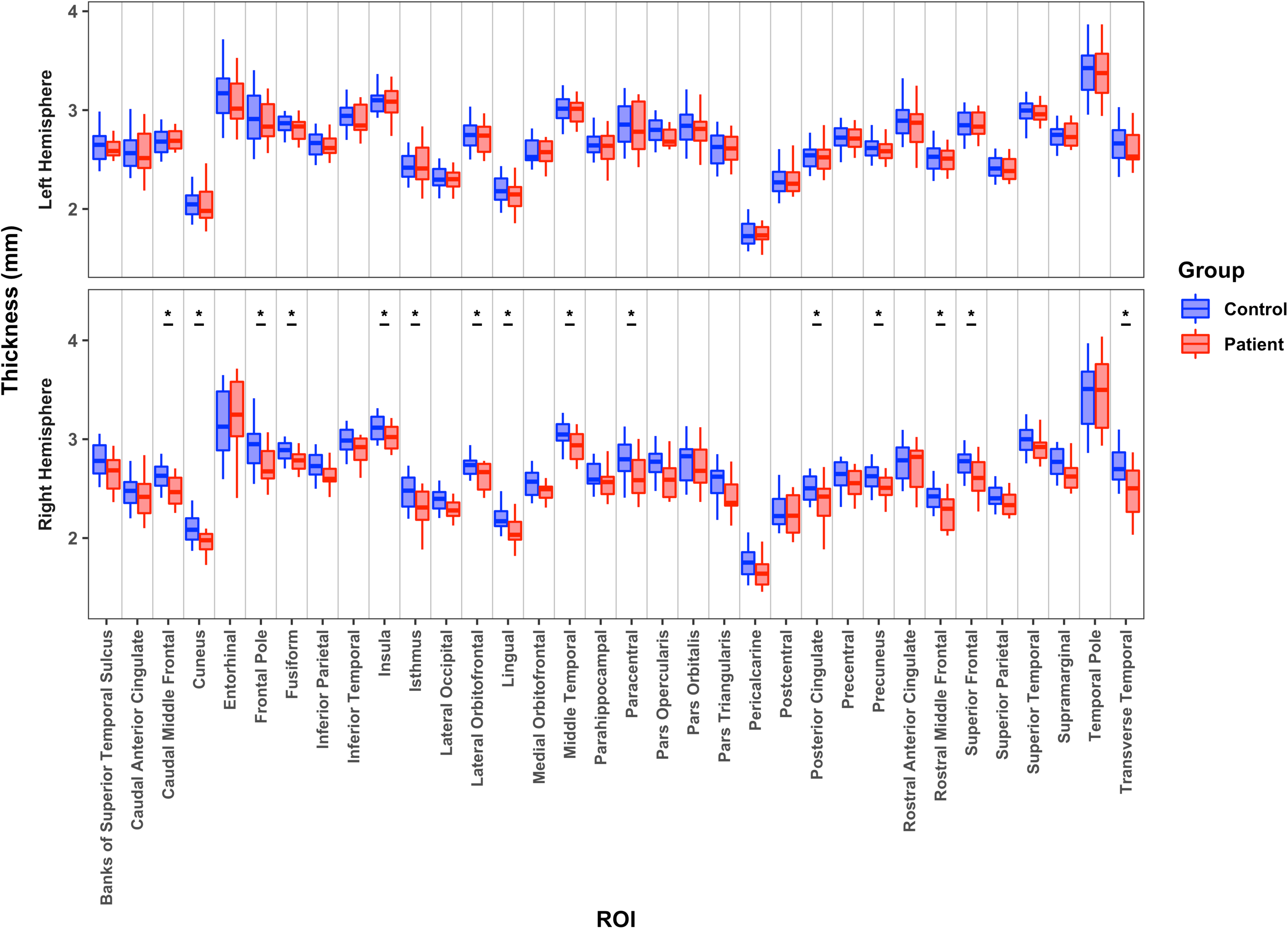
Median cortical thickness for 34 cortical regions for patients with a preserved left hemisphere or right hemisphere relative to their matched controls. *: *p*<0.05; **: *p*<0.01; ***: *p*<0.001.

A forward regression analysis with CxT indices of all 34 regions of LH patients vs. LH controls resulted in no predictors passing initial criterion for inclusion, indicating that we cannot differentiate patients from controls. A model with a very strong association between the predictors and RH patient vs. RH control group, *r^2^*=0.8, included the CxT of the caudal middle frontal, frontal pole, lateral occipital, lingual, pars orbitalis, and transverse temporal regions. A model with very strong prediction, *r^2^*=0.89, which included CxT of entorhinal, rostral middle frontal, and superior temporal regions differentiated LH from RH patients. No viable model was able to separate the patients into high vs. low binned ILAE outcome scores.

#### 3.2.2. Cortical Surface Area

As depicted in Figure 3, compared with their matched controls, patients with a preserved LH did not differ on CSA, whereas patients with a preserved RH had significantly greater CSA in lateral orbitofrontal, paracentral, and parahippocampal cortices. The two patient groups differed from one another with the preserved LH patients having greater CSA in three regions (pars opercularis, rostral anterior cingulate and transverse temporal) and the preserved RH patients having greater CSA in four regions (frontal pole, inferior parietal, parahippocampal and pars orbitalis). (See Supplementary Table 3 for individual *p* and *BF* values.)

**Figure 3:**
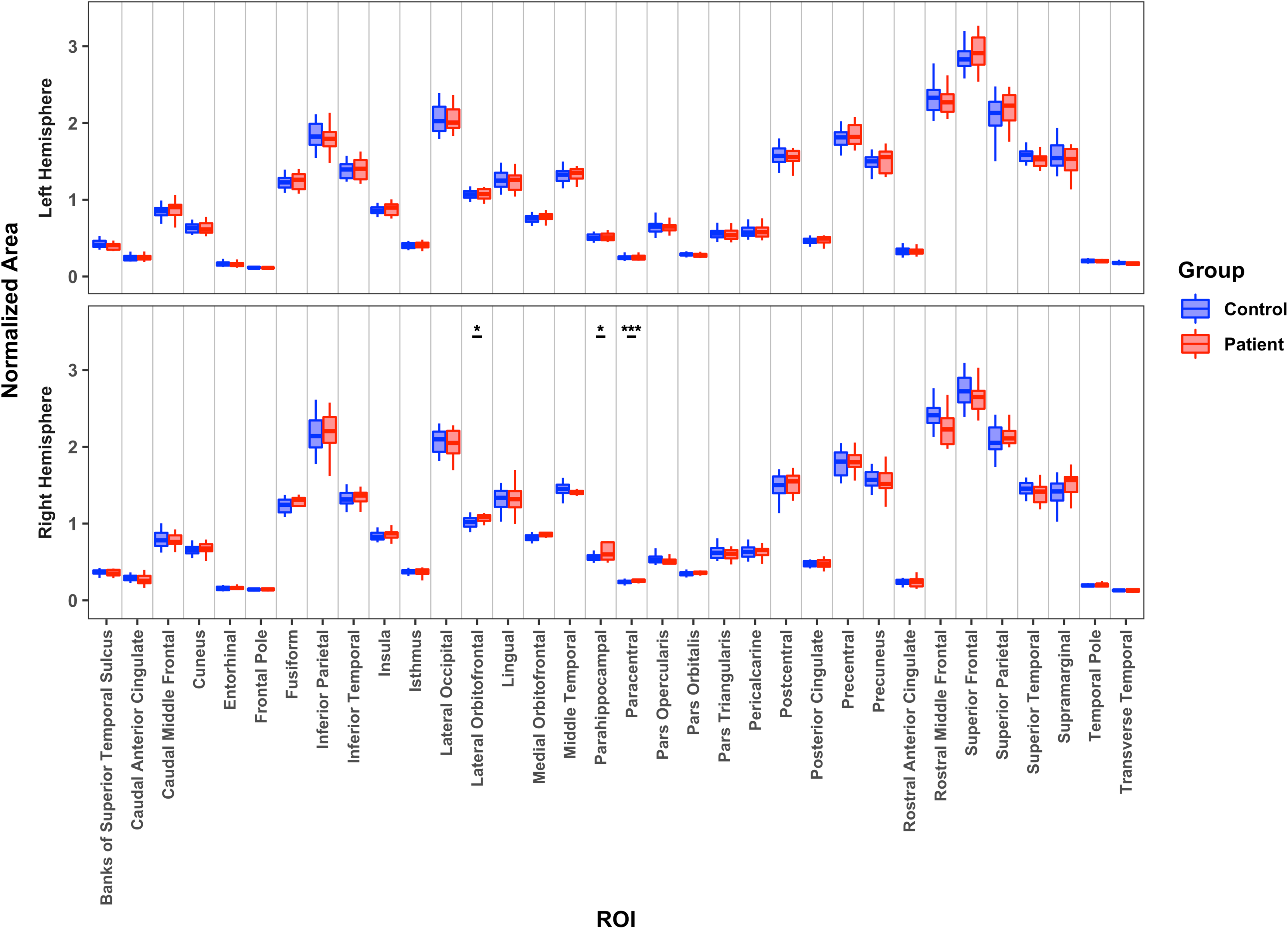
Median normalized cortical surface area for 34 cortical regions for patients with a preserved left hemisphere or right hemisphere relative to their matched controls. *: *p*<0.05; ***: *p*<0.001.

In a forward logistic regression analysis with CSA of all 34 areas entered as independent measures for the LH patients and matched controls, we could not derive a model associated predictor (region) with outcome (group membership). For the RH patients and controls, the most predictive model with a very strong association value of *r^2^*=0.83 included the CSA of the entorhinal, lateral orbitofrontal, paracentral, rostral middle frontal, and superior temporal regions. The two patient groups were differentiable by a model containing only the CSA of the pars orbitalis area, with a very strong prediction, *r^2^*=0.95, and a model with moderate prediction, *r^2^*=0.33, including only the paracentral area predicted binned ILAE outcome score.

#### 3.3.3. Cortical Volume

On the final dependent measure, CV, as shown in Figure 4, the GLM revealed no differences between the LH patients and the LH of the controls. A difference in rostral middle frontal volume was observed between the RH patients and the RH of the controls. A direct comparison between the two patient groups indicated four regions with less volume in the RH than the LH patient group (inferior parietal, pars opercularis, transverse temporal, rostral anterior cingulate) and greater volume in the preserved RH than LH patient group in two regions (pars orbitalis, superior frontal). (See Supplementary Table 4 for individual *p* and *BF* values.)

**Figure 4:**
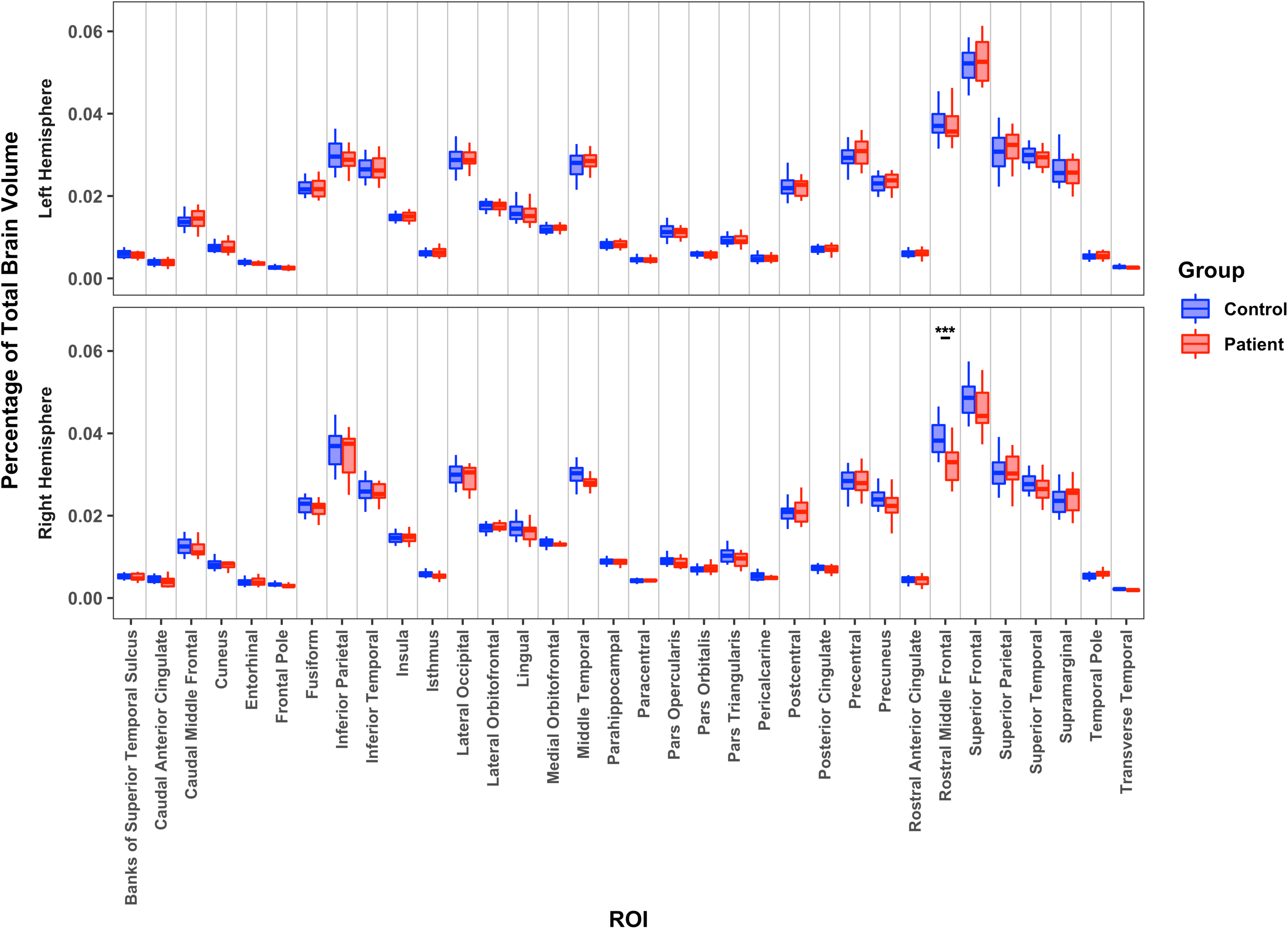
Median region volume (normalized by total volume) for each of 34 cortical regions for patients with a preserved left hemisphere or right hemisphere relative to their matched controls. ***: *p*<0.001.

A forward logistic regression analysis with the measures of CV for each of the 34 regions predicting the differences between LH patients and controls had no significant predictors. A model with strong predictive value, *r*^2^=0.57, which included the rostral middle frontal and pars orbitalis volume predicted group membership for the RH patients vs. RH controls. The RH and LH patients were well differentiated by a model containing pars opercularis and pars orbitalis with very strong predictive value, *r*^2^=0.9. Additionally, a model containing CV of paracentral, pars triangularis, and superior parietal volume strongly predicted ILAE score, with *r*^2^=0.71.

In summary, the morphometric differences were more pronounced for the comparison of the RH patient group and the corresponding RH of the controls than for the LH patient/control comparison and this was true across all three measures and modes of analysis (GLM, bivariate regression). As shown in the lefthand column of Figure 5, the differences in the patients’ preserved RH appear to be largely in language-related regions including pars orbitalis, which borders on Broca’s area, as well as superior temporal and additional frontal regions. We also observe RH patients’ differences relative to controls in homologous language-related regions, in the comparison of the two patient groups against each other with differences in the superior temporal, pars orbitalis, and pars opercularis (area 44 and part of Broca’s area).

**Figure 5:**
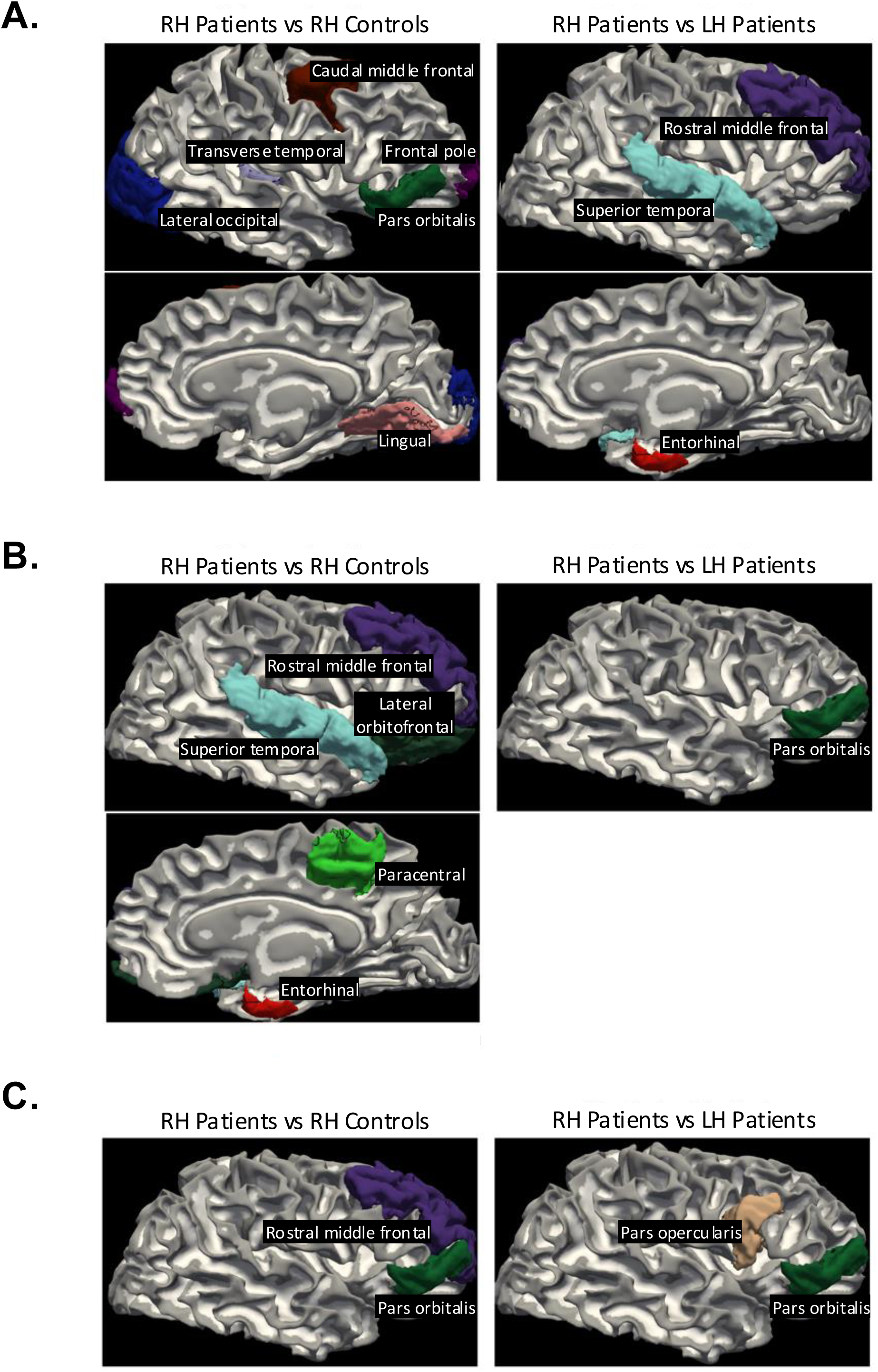
Regions that significantly distinguished between groups (right hemisphere patients versus controls in left columns, right hemisphere patients versus left hemisphere patients in right columns) per binary logistic regression modelling for (A) cortical thickness, (B) cortical surface area, and (C) cortical volume.

### 3.4. Subcortical Regional Morphometrics

We segmented (using FreeSurfer) the preserved hemisphere in patients and both hemispheres in controls into nine subcortical regions, namely, the nucleus accumbens (here referred to as accumbens), amygdala, caudate, cerebellum, hippocampus, pallidum, putamen, thalamus, and ventral diencephalon. We extracted the volume of these regions.

Compared with the LH of controls, patients with a preserved LH showed a significantly lower percentage of whole brain volume of the accumbens, caudate, pallidum, and putamen. Patients with a preserved RH had significantly less volume in the accumbens and hippocampus compared to controls. In the direct contrast between the two patient groups, the LH patient group had significantly less volume in the caudate and putamen relative to the RH patient group (Figure 6). (See Supplementary Table 5 for individual *p* and *BF* values.)

**Figure 6:**
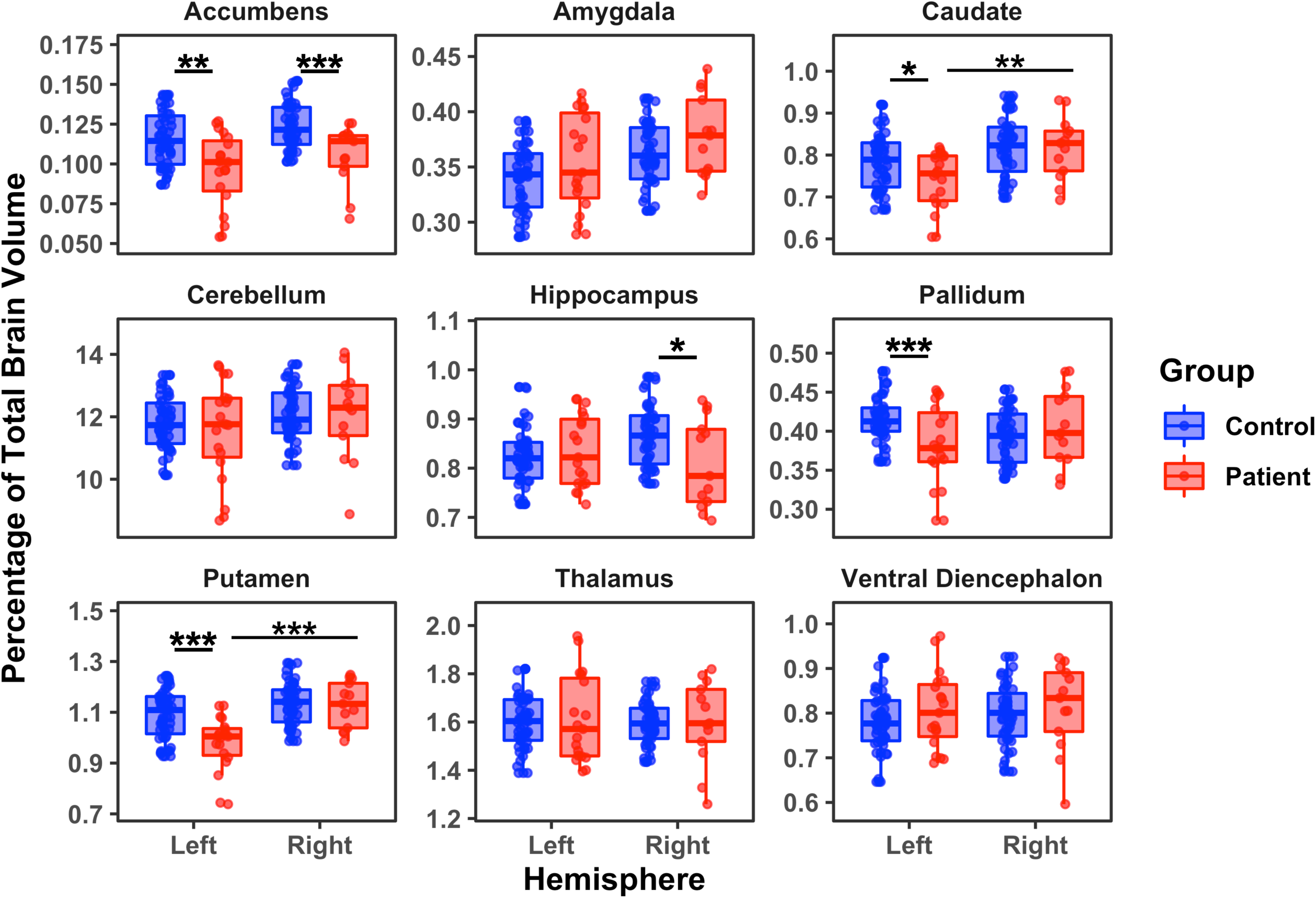
Subcortical regional volumes (normalized) for patients with a preserved left hemisphere or right hemisphere relative to their matched controls *: *p*<0.05; **: *p*<0.01; ***: *p*<0.001.

To evaluate which of these nine subcortical measures, singly or in combination, best predicted group membership, we used the same forward binary logistic regression procedure as above. The model differentiating LH patients and LH controls yielded a model which included putamen and accumbens, with *r^2^*=0.44, revealing a strong relationship between these two predictors and group membership. The same analysis for RH patients and RH controls included only the accumbens and yielded an *r^2^*=0.28, indicating a moderate relationship between this predictor and outcome. A regression model that included the putamen volume alone differentiated strongly, *r^2^*=0.52, between the preserved RH vs. LH patients. Last, no model including volume of the subcortical measures was reliable in classifying patients into the two ILAE outcome scale bins.

In sum, the GLM results clearly differentiated the LH and RH patients from their matched controls on four and two of the nine subcortical regions, respectively, with the accumbens having lower volume for both patient groups vs. controls. The preserved LH patients had less volume than the RH patients in the putamen and caudate. The regression model identified the accumbens and the putamen as predicting LH patients from controls and the accumbens alone as predictive of group membership in RH patients vs. controls. We could not predict the ILAE outcome scale by any variable/s. These findings indicate that the accumbens and/or putamen are key subcortical structures that differentiate patients from controls, and the putamen plays a vital role in differentiating the side of preserved hemisphere in the patients.

### 3.5. Analysis Excluding Ablation Cases

One possible explanation for the differences between the findings for the LH and RH patients is that the number of cases of resection vs. ablation vary for the two groups. More LH than RH patients have an ablation, which implicates a much smaller region of anatomical change compared to a resection (LH – 7 ablation, 12 resection; RH – 2 ablation, 11 resection). To ensure that the results reported above are not a consequence of this imbalance, we recomputed all the binary logistic regression analyses with only the resection cases (12 LH, 11 RH). Notwithstanding the loss of statistical power with this data reduction, the findings using the three gross measures (LV, GM, and WM) largely mirrored that of the analysis including all patients: whereas no model could predict group membership between LH patients and controls or between LH patients and RH patients, a model with moderate predictive power, *r*^2^ = 0.48, with LV and GM discriminating between the RH patients and RH controls.

Additionally, the analysis of the cortical regions with only the resection patients revealed largely similar results to that conducted with all the patients. With respect to CxT, a model with weak to moderate predictive value, *r*^2^=0.27 – which included the fusiform and inferior parietal regions – predicted group membership for LH patients vs. LH controls. We observed a strong association between group membership (patient vs. control) for the RH with the CxT of only two areas, the frontal pole and posterior cingulate, playing a predictive role, *r*^2^=0.43. We also obtained a strong prediction of side of preserved hemisphere (LH vs. RH) for separating the two patient groups, *r^2^*=0.93 based on differences in pars orbitalis CxT. For CSA, no model predicted patients vs. controls for the LH comparisons. However, a model with strong predictive power (including lateral orbitofrontal, parahippocampal, rostral middle frontal, superior temporal regions), *r*^2^=0.76, separated patients and controls in the RH comparisons, and a very strong model of *r*^2^=0.90, only with the pars orbitalis separating the two patient groups. For CV, whereas no model was statistically viable for separating the controls and LH patients, a model with two variables − rostral middle frontal region and pars orbitalis − segregated the controls and RH patients with strong predictive power, *r*^2^ =0.53. A model with a single variable, pars opercularis, was strongly predictive of whether patients had the LH or RH preserved, *r*^2^=0.69. Lastly, no model of the cortical measures was reliable in classifying the LH and RH patients into the two ILAE outcome scale bins.

Finally, the regression analysis using the subcortical regions largely replicated the analysis with all the patients. A model with strong association, *r*^2^=0.56, that included the accumbens and putamen separated the LH patients from their controls; a reasonably predictive model with *r*^2^=0.33 and including the accumbens separated the RH patients from their controls, while a model with strong association, *r*^2^=0.66, which included only the putamen, separated the two patient groups.

That the findings are almost all replicated with a subset of only those patients who have resections indicates that the results when we included all patients are not an obvious artifact of differences in post-surgical anatomical size of resection/ablation.

### 3.6. Correlations Between Gross Morphometrics and Cortical/Subcortical Morphometrics

In this final analysis, we examined whether any of the gross measures − for example, the size of the LVs − is statistically associated with cortical or subcortical measures. Because the cranium is a fixed size cavity, a large increase in one major feature/tissue such as LV volume will necessarily induce a large decrease in the remaining WM (Lopez et al., 2022). To examine this, we performed bivariate correlations (with multiple comparison correction *p*<.0002) between the gross measures and each of the volumes of subcortical regions and then between the gross measures and each of the three dependent measures for each of the 34 cortical parcels. We did this with data from all patients and then separately for LH patients and RH patients. In no analysis did any correlation survive the most stringent multiple comparison, suggesting that group differences in gross anatomy cannot be explained simply by any differences in cortical or subcortical volume.

## 4. Discussion

We undertook a comprehensive characterization of the structural integrity of the post-surgical preserved hemisphere in DRE pediatric patients, including the evaluation of gross anatomy, CxT, CV, and CSA of 34 cortical parcels, as well as the volume of nine subcortical regions. Studies of post-surgical brain morphometry conducted with adults with DRE have yielded conflicting findings: whereas some have reported reduction, and even reversal, of progressive cortical thinning (Federico & Wiebe, 2020; Galovic et al., 2020; Galovic et al., 2019; Li et al., 2022; McDonald, 2021) and CV and atrophy (Federico & Wiebe, 2020), others have noted ongoing, progressive atrophy (and other postsurgical changes (Janson et al., 2024)) even in regions remote from the resection site (Caciagli et al., 2017; Park et al., 2022). Almost all such studies have been carried out in adults − primarily those with temporal lobe epilepsy and anterior temporal lobectomy − and have largely focused on just one variable, such as CxT or CV (but see (Zhao et al., 2021) for both).

A clear depiction of the non-resected hemisphere is also important in pediatric cases and is increasingly pressing as the anatomical integrity of the preserved hemisphere is linked with post-surgical behavioral function in this population (Skirrow et al., 2015; Zhao et al., 2021). Furthermore, surgery for those with DRE, especially during childhood, is becoming a first-line form of intervention (Consales et al., 2021; Cross et al., 2022; Vakharia et al., 2018), which can resulting in cessation of the downward presurgical neuropsychological trajectory (Eriksson et al., 2024). It is also unwise to extrapolate findings from the studies of adult patients to pediatric patients given the greater potential for plasticity in the latter over the former.

Here, we recruited 32 post-surgical patients, with resections ranging from large removal of tissue, as in cases with hemispherectomy, to more focal alterations such as after robot-assisted laser ablation. On all metrics, we first compared the patients’ preserved hemisphere to the corresponding hemisphere of the matched controls and then compared directly the morphometry of the patients with a preserved LH with that of patients with a preserved RH. In addition to the detailed measurement of gross anatomical indices and integrity of cortical and subcortical regions, we also examined differences as a function of which hemisphere was preserved and analyzed correlations across the various metrics. Last, to ensure that the findings were not an artifact of whether the surgical intervention was a resection versus an ablation, we recomputed all analyses while excluding those patients with laser ablation. All statistical analyses were conducted using both GLM and binary logistic regression analyses, and the results were largely, although not always perfectly, converging. Several key findings emerged from the data (see summary in Table 2).

**Table 2.**
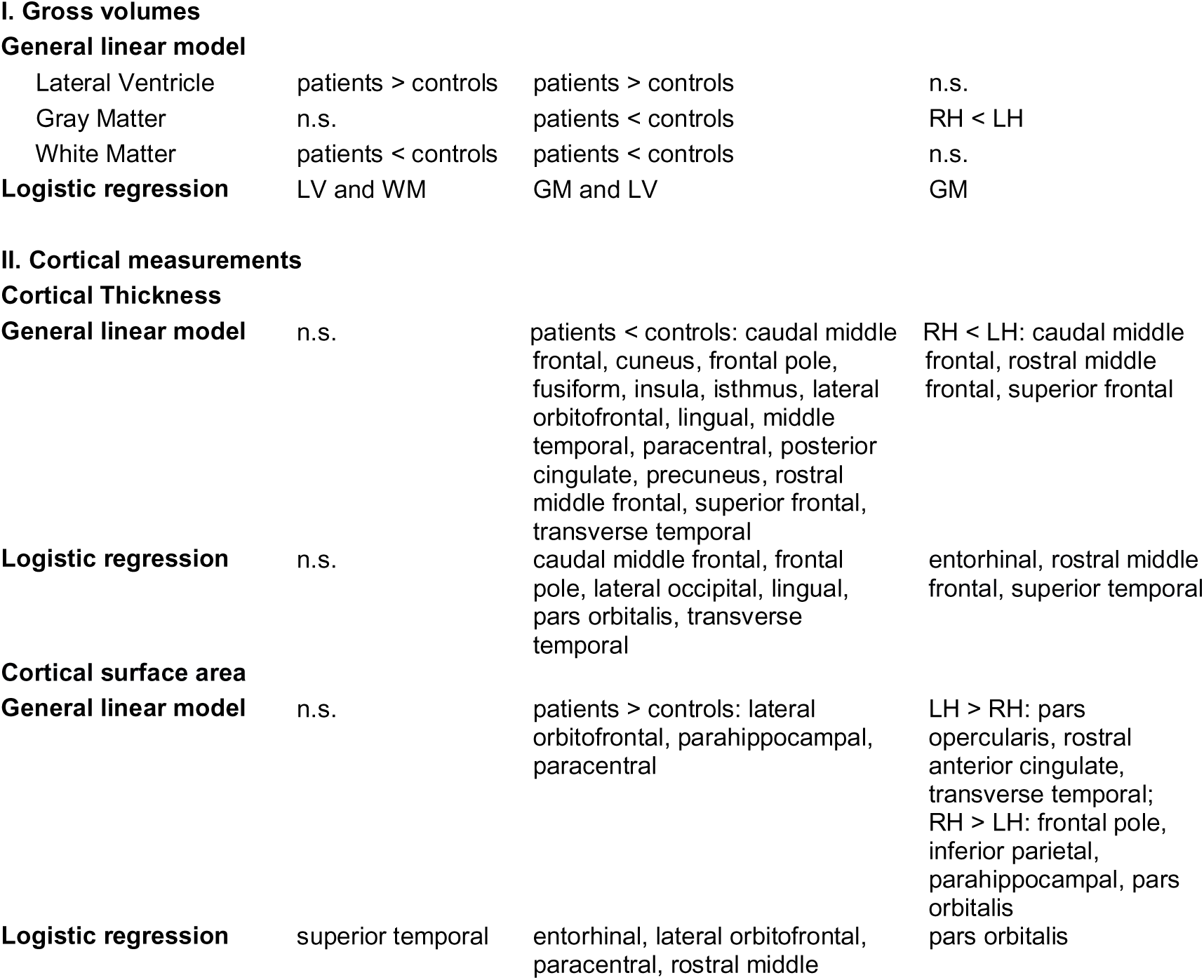
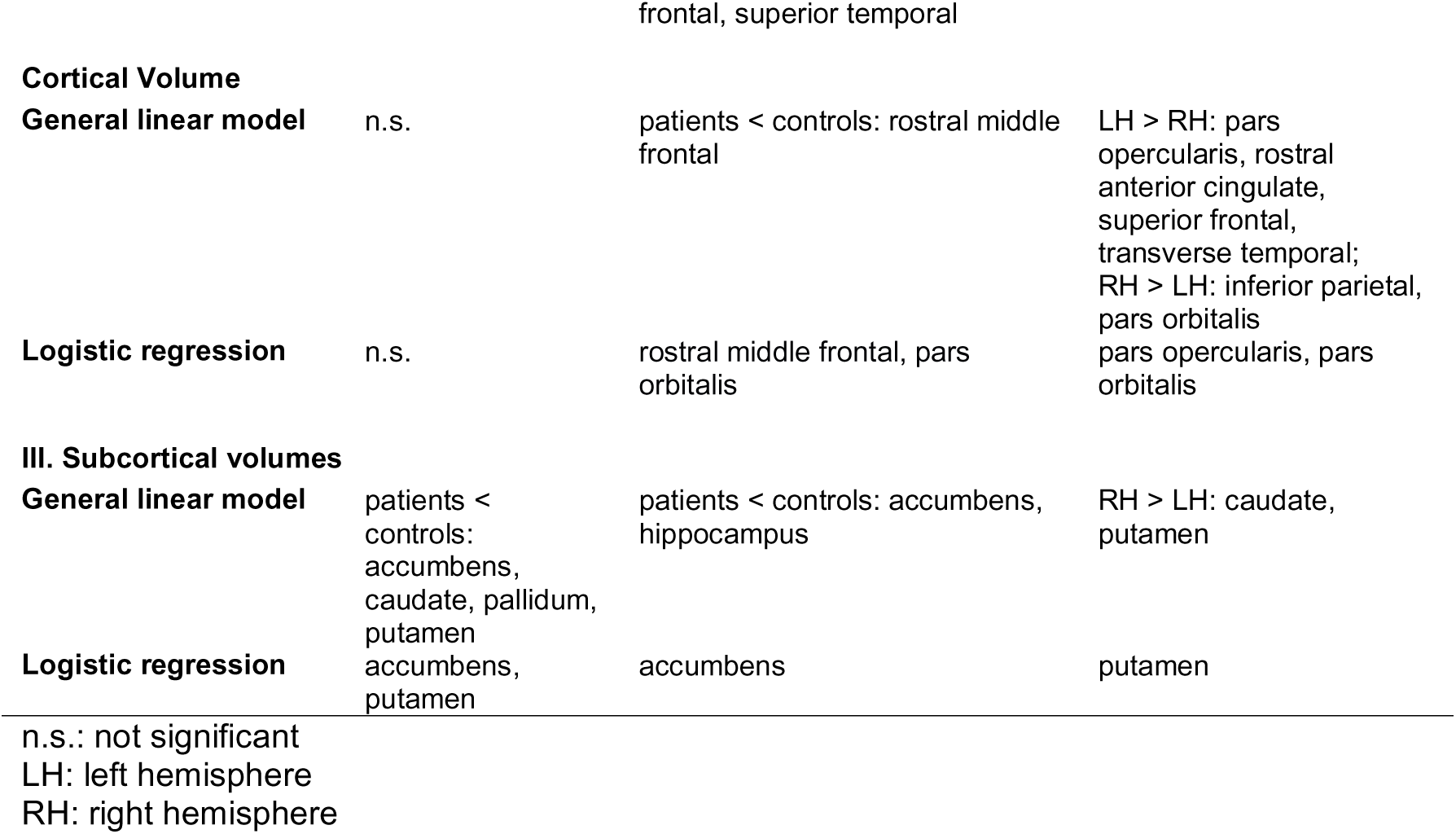
At-a-glance summary of all results for univariate and multivariate analyses.

First, both LH and RH patient groups differed from their matched controls, having larger LVs and less WM; only RH patients had less GM than controls. The volume of GM alone successfully predicted whether patients had a preserved LH vs. RH with the former having less atrophy than the latter. In the analysis of the 34 cortical regions, the morphometric measures of the brains of patients with a preserved LH were more like the corresponding hemisphere of controls than was the case for patients with a preserved RH. The preserved RH patients differed from controls on multiple dependent measures in both the GLM analyses and binary logistic regressions. For example, we observed lower CxT in a host of frontal, temporal, and occipital areas and in the isthmus, insula, and precuneus. These patients also had greater CSA than controls in lateral orbitofrontal, parahippocampal, and paracentral areas, and a subset of these areas predicted the group membership (patient vs. control) in the best fit regression model. Lastly, the RH patients had less CV of the rostral middle frontal area (and no obvious parietal region differences) relative to controls, which, together with the pars orbitalis gave rise to appropriate assignment of membership into patients vs. controls. The striking contrast between the LH vs. RH preserved patients is clear, as RH but not LH patients consistently showed morphometric differences from controls.

By inference, unsurprisingly, the direct comparison between the two patient groups also revealed a host of differences for preserved RH vs. LH, with lower CxT in frontal, entorhinal, and superior temporal regions, less CSA in frontal cingulate and transverse temporal regions, but also greater CSA in, for example, frontal pole, parahippocampal and pars orbitalis. Lastly, those with a preserved RH had less CV (more atrophy) in a set of frontal and temporal regions, and, consistent with the CSA measurements, greater CV in the inferior parietal and pars orbitalis regions. Between patient groups, differences in the pars orbitalis and the pars opercularis manifested as key sites in the CSA and CV in both the GLM and regressions analyses.

With respect to the nine subcortical regions, the volume of the putamen and accumbens (along with other structures such as pallidum, caudate, and hippocampus) was lower in both patient groups vs. controls, and the volume of the putamen alone (less volume in LH than RH) was able to differentiate the two patient groups from each other. The indices for the gross measures showed no correlation with any other subcortical or cortical metrics that survived stringent familywise correction, indicating independence between the morphometric changes of LV, WM and GM volume, and other measurements and confirming the findings that morphometric differences may be more region-specific than global (Sele et al., 2021).

These findings are consistent with some results in the literature in which, relative to controls, children with focal epilepsy showed differences in CxT in the frontal lobe and to a greater degree in the RH than LH and less CV (Taherahmadi et al., 2025) (also see (Fonseca et al., 2023)). Another study with children showed extensive cortical thinning predominantly in frontal and temporal regions (Li et al., 2020), as did a third study which also documented changes in frontal and temporal regions (but also in occipital cortex in children) (mean age ∼10.5 years) (Garcia-Ramos et al., 2015); the latter study also showed, as we did, changes in the putamen.

Machine learning algorithms have also been successful in classifying structural brain properties into two groups, one with epilepsy and the other healthy controls. Interestingly, out of the most commonly used measurements which include CxT, GM volume, WM volume, and cerebrospinal fluid volume, the GM and WM volumes were most informative, accounting for 80% of the data (Azzony et al., 2023). We too observed changes in gross anatomical measurements, but also in the size of the LVs, and such changes were present to a greater degree after LH than RH resection. Azzony et al. (2023) also applied different machine learning classifiers to data extracted from 100 ROIs with K-nearest neighbor approach yielding 97% accuracy in separating control vs. patient group membership. Our logistic regression procedure, while not always as accurate as those of Azzony et al. (2023), is able to classify our patient vs. control groups, especially for RH preserved patients and their controls and did so using only a small number of datapoints (34 for cortical, 9 for subcortical and 3 for gross anatomy).

Especially notable in the contrast between the previous findings and the current study is that the studies reviewed here examined morphological differences in patients with childhood epilepsy, but no patient had undergone surgery. That differences are evident in those with focal epilepsy but without resection suggests that the altered morphometry we document is more likely a consequence of the epileptic activity than an outcome of the surgery per se. Last, these findings may be specific to the epilepsy population as they are not observed in other neurological disorders in childhood. Children with traumatic brain injury do not show differences in total brain, WM, or GM volumes or regional subcortical volumes, although cortical thinning was noted in the angular gyrus and basal forebrain as well as in frontal and occipital regions shortly after injury (but this resolved over several months) (Ware et al., 2023).

The key findings we present here include numerous differences in all three dependent measures between those with a preserved RH and their matched controls or between them and those with a preserved LH. There are far fewer differences in those with a preserved LH and their matched controls. In almost all instances, the preserved RH has reductions in CxT and CV but, surprisingly, has larger CSA in a few regions including orbitofrontal and parahippocampal regions, along with alterations in subcortical regions, namely in the accumbens, caudate, and putamen.

Because we matched the two groups on age, gender, age at first surgery, age of seizure-onset age, and ILAE seizure outcome scores, and we included age and gender as covariates in all GLM analyses, none of these factors obviously account for the hemispheric differences. One potential difference is that the data were acquired on two different scanners, with arguably different resolution, but this, too, does not obviously explain the asymmetry as both patient groups were roughly equally represented on each scanner (Siemens Verio: 4 RH, 9 LH and Siemens Prisma: 8 RH, 8 LH) and, moreover, the data were harmonized across magnets. We have also ruled out another potential confound concerning a possible difference in the size of the surgery in each group: the re-analysis of the data with only the frank resection cases and no ablation patients, rules out this potential confound. In fact, despite the reduction in statistical power by removing the ablation patients, we replicate almost all the results of the analyses with the data including all the patients. The asymmetry for the preserved LH and RH is also not obviously explained by differences in lobar resections as the LH and RH groups with more focal resections had a roughly equal number of patients with temporal (LH: 4, RH: 4), frontal (LH: 2, RH: 2), occipital (LH: 1, RH: 2), and parietal (LH: 1, RH: 1) resections.

A key to understanding the hemispheric differences comes from scrutinizing the regional differences between those with preserved LH vs. RH. Of note, almost all the areas that differ in the preserved RH vs. LH or RH vs. controls are homotopic with or proximal to standard LH language areas, for example, the caudal middle and superior frontal, middle and superior temporal, and transverse temporal regions (see Figure 5). These findings, derived from an unbiased and whole brain data-driven analysis appear to converge largely on regions that are implicated in language function. Resecting some, most, or all the LH (preserved RH group), which is likely the native dominant language areas in most of the patients, may be the trigger for alterations of LH homologues of language areas in the preserved RH. In contrast, a resection of some, most, or all the RH (preserved LH group) results in remarkably few structural or anatomical differences relative to the matched controls’ LH. Our results are also consistent with one study that revealed that, after left arcuate fasciculus pediatric resection, an increase in the right homotopic region was evident (Goradia et al., 2011). Although other studies have shown widespread changes in structural connectivity in the contralateral hemisphere (for example, (Jeong et al., 2016)), our findings have uncovered specificity in the direction (only in those with LH resection) and region (largely language-related areas) of post-surgical structural alterations.

In summary, with LH cortical resection, regions of the preserved RH that are homotopic with LH language regions are structurally different from controls’ RH. The process of accommodating language function in the RH, which likely has some nascent or “shadow” of language function (Martin, Seydell-Greenwald, Berl, Gaillard, Turkeltaub, et al., 2022; Newport et al., 2022)), potentially underlies the morphometric changes in the RH. Much evidence attests to the fact that, in early childhood, both hemispheres are predisposed to language function (Bates et al., 2001), and even adolescents and young adults who suffer a perinatal ischemic stroke to the LH nevertheless display sentence processing abilities equal to that of non-neurological controls, presumably a result of RH language engagement (Martin, Seydell-Greenwald, Berl, Gaillard, & Newport, 2022; Newport et al., 2022). The idea of RH plasticity or upregulation for language has also been confirmed in neuroimaging studies in which, for example, during a silent word generation, individuals with peri- or prenatal periventricular LH damage evince RH blood-oxygen-level-dependent activation task equal to that of the LH in right-handed controls (Staudt et al., 2002). Findings such as these attest to the remarkable plasticity and potential for functional reorganization and/or upregulation of language to the RH homotopic frontotemporal regions.

What remains unexplained is why we see, relative to controls, a reduction in the morphometric measures of the RH homotopic language regions, rather than a maintenance or even expansion, which is often associated with the assumption of a new function. Presumably, as is also true over the course of typical development, cortical regions are pruned as functions are acquired through the removal of inefficient synapses, dendrites, and neurons (Bourgeois & Rakic, 1996; Changeux & Danchin, 1976; Uddin et al., 2015). Many have suggested that the cortical tissue loss reflects improved neural processing by winnowing brain circuits for select functions (Natu et al., 2019). Cortical thinning is coupled with morphological changes (Natu et al., 2019) and, in childhood, GM thinning specifically is associated with changes in behavior (Sowell et al., 2004), which may be associated with increases in WM (Giedd et al., 1999). Relevantly for this paper, in a longitudinal study, changes of CxT asymmetry in the triangular part of the IFG, which we also identified here, and constituting a portion of Broca’s area in the LH, is suggestive of a neural correlate of language improvement in children’s brains at roughly ages 5-7 yr (Qi et al., 2019).

Notably, our available data sample consisted of primarily postoperative scans. Given the cross-sectional, rather than longitudinal design, of the current study, we have been unable to evaluate changes over time and their relationship to behavior. Because of our one-time snapshot, whether the profiles of the childhood DRE brains changed relative to controls’ brains specifically over the pre-surgical period and the relationship between these changes over time, hemisphere of resection, and cognitive (specifically language) outcome remain unaddressed and serve as rich fodder for future studies. Specifically, reversal of progressive thinning and recovery from atrophy and thinning (Zhao et al., 2021), which is associated with improved language function (Li et al., 2025) − and specifically, in post-surgical vs. pre-surgical cortical regions − are promising targets for future study and may add to efforts to differentiate and predict the three possible outcomes, as laid out by Sainburg et al (2025), namely, damage and degeneration, recovery, and reorganization.

## CRediT authorship contribution statement

Michael Granovetter: Conceptualization, Methodology, Software, Validation, Formal analysis, Investigation, Data curation, Writing - Review & Editing, Visualization, Project administration. Marge Maallo: Conceptualization, Methodology, Software, Validation, Formal analysis, Investigation, Data curation, Writing - Review & Editing, Visualization, Project administration. Christina Patterson: Resources, Writing - Review & Editing. Daniel Glen: Methodology, Software, Writing - Review & Editing. Marlene Behrmann: Conceptualization, Methodology, Formal analysis, Writing - Original draft, Supervision, Project administration, Funding acquisition.

## Declaration of generative AI in scientific writing

Generative AI has not been used in the writing of this paper.

## Acknowledgment and funding sources

We are grateful to our study participants, including patients, the controls, and their families, who generously donated their time to make this research possible. We also thank the Pediatric Epilepsy Surgery Alliance, especially Monika Jones, JD, Chief Executive Officer. We are grateful to Scott Kurdilla, Dr. Tina Liu, Sophia Robert, Mark Vignone, and Debbie Viszlay for assistance with data acquisition; and Dr. Vladislav Ayzenberg, Dr. Nicholas Blauch, Dr. Alison Butler, Dr. Samuel Dienel, Dr. Joel Greenhouse, Dr. Alexander Layden, Dr. Timothy Verstynen, and the CMU Visual Cognition group for their helpful discussions and feedback.

This research was supported by Award Numbers T32GM081760 and T32GM144300 from the National Institute of General Medical Sciences (M.C.G.), R01EY027018 from the National Eye Institute (M.B. and C.P.), a fellowship from the American Epilepsy Society (#847556) (M.C.G.), a scholarship from the American Psychological Foundation (M.C.G.), and funding from the CMU-Pitt BRIDGE Center. DRG was supported by the National Institute of Mental Health Intramural Research Program (ZICMH002888) of the National Institutes of Health/Health and Human Services, USA. M.B. also acknowledges support from P30 CORE award EY08098 from the NEI, NIH, and unrestricted supporting funds from The Research to Prevent Blindness Inc, NY, and the Eye & Ear Foundation of Pittsburgh. The content is solely the responsibility of the authors and does not necessarily represent the official views of NIGMS, NEI, NIMH, AES, or APF.

## Declaration of competing interest

The authors of this manuscript declare hereby that they have thoroughly reviewed and evaluated potential competing interests related to the work reported in this paper. Marlene Behrmann is a founder of Precision Neuroscopics. Aside from this, after careful consideration, it has been determined that there are no known competing financial interests, personal relationships, or other factors that could influence the integrity or objectivity of the presented research. The authors are committed to upholding the highest standards of transparency and academic honesty in their work.

## Data availability

A DOI has been reserved on Carnegie Mellon University’s KiltHub repository (operated via Figshare): 10.1184/R1/24153423. All relevant data and code will be published in this repository upon publication of the article.

## Supplementary material

Supplementary material is available online.

## Notes

### Summary of Updates

Text and figures of the manuscript have been updated.

