## Supplementary Material for "Morphometrics of the preserved post-surgical hemisphere in pediatric drug-resistant epilepsy and implications for post-operative cognition"

**Table S1 R packages**

| Package | Version | Use |
|---|---|---|
| car[1] | 3.0-11 | Summarizing statistics |
| DescTools[2] | 0.99.44 | Data quality control |
| lme4[3] | 1.1-27.1 | Fitting linear mixed models |
| neuroCombat[4] | 1.0.9 | Harmonization |
| psych[5] | 2.1.9 | Computing descriptive statistics |
| stringr[6] | 1.4.0 | Data tidying |
| tidyverse[7] | 1.3.1 | Data tidying |

**Table S2 Descriptive statistics: Median (median absolute deviation) of gross morphometrics**

| | Control LH | Control RH | Patient LH | Patient RH |
|---|---|---|---|---|
| LV | 5.36x10^3 (1.67x10^3) | 4.97x10^3 (1.60x10^3) | 6.33x10^3 (3.52x10^3) | 5.65x10^3 (3.81x10^3) |
| GM | 2.83x10^5 (2.59x10^4) | 2.84x10^5 (2.32x10^4) | 2.68x10^5 (3.03x10^4) | 2.44x10^5 (2.77x10^4) |
| WM | 2.17x10^5 (2.87x10^4) | 2.16x10^5 (2.92x10^4) | 2.08x10^5 (2.53x10^4) | 1.99x10^5 (2.56x10^4) |

LH = left hemisphere
RH = right hemisphere
LV = lateral ventricle
GM = gray matter
WM = white matter

**Table S3 Descriptive statistics: Median cortical thickness (Median absolute deviation) per cortical region**

| Region | Control LH | Control RH | Patient LH | Patient RH |
|---|---|---|---|---|
| Banks of Superior Temporal Sulcus | 2.65 (1.92x10^-1) | 2.78 (2.17x10^-1) | 2.59 (9.72x10^-2) | 2.68 (2.73x10^-1) |
| Caudal Anterior Cingulate | 2.56 (1.98x10^-1) | 2.48 (1.54x10^-1) | 2.51 (3.12x10^-1) | 2.42 (2.45x10^-1) |
| Caudal Middle Frontal | 2.68 (1.57x10^-1) | 2.63 (1.38x10^-1) | 2.69 (1.35x10^-1) | 2.46 (1.92x10^-1) |
| Cuneus | 2.05 (1.46x10^-1) | 2.08 (1.72x10^-1) | 1.98 (1.38x10^-1) | 1.98 (1.33x10^-1) |
| Entorhinal | 3.17 (2.80x10^-1) | 3.13 (4.66x10^-1) | 3.01 (2.36x10^-1) | 3.25 (4.93x10^-1) |
| Frontal Pole | 2.91 (3.22x10^-1) | 2.95 (1.96x10^-1) | 2.83 (1.76x10^-1) | 2.67 (3.05x10^-1) |
| Fusiform | 2.87 (1.02x10^-1) | 2.89 (1.07x10^-1) | 2.83 (1.46x10^-1) | 2.79 (9.55x10^-2) |
| Inferior Parietal | 2.67 (1.43x10^-1) | 2.73 (1.63x10^-1) | 2.62 (1.25x10^-1) | 2.6 (1.00x10^-1) |
| Inferior Temporal | 2.94 (1.34x10^-1) | 2.99 (1.61x10^-1) | 2.84 (1.42x10^-1) | 2.92 (1.38x10^-1) |
| Insula | 3.1 (1.52x10^-1) | 3.12 (1.72x10^-1) | 3.09 (1.79x10^-1) | 3.02 (1.71x10^-1) |
| Isthmus | 2.42 (1.58x10^-1) | 2.48 (2.05x10^-1) | 2.41 (1.99x10^-1) | 2.31 (2.12x10^-1) |
| Lateral Occipital | 2.3 (1.39x10^-1) | 2.4 (1.17x10^-1) | 2.3 (1.10x10^-1) | 2.28 (1.19x10^-1) |
| Lateral Orbitofrontal | 2.75 (1.59x10^-1) | 2.74 (1.09x10^-1) | 2.74 (1.78x10^-1) | 2.67 (1.58x10^-1) |
| Lingual | 2.18 (1.37x10^-1) | 2.17 (8.76x10^-2) | 2.15 (1.69x10^-1) | 2.03 (1.75x10^-1) |
| Medial Orbitofrontal | 2.53 (1.21x10^-1) | 2.57 (1.85x10^-1) | 2.57 (1.80x10^-1) | 2.49 (7.38x10^-2) |
| Middle Temporal | 3.01 (1.43x10^-1) | 3.05 (1.39x10^-1) | 3.01 (1.24x10^-1) | 2.94 (1.99x10^-1) |
| Parahippocampal | 2.64 (1.27x10^-1) | 2.59 (2.02x10^-1) | 2.64 (1.65x10^-1) | 2.57 (1.38x10^-1) |
| Paracentral | 2.85 (2.67x10^-1) | 2.8 (1.99x10^-1) | 2.78 (3.91x10^-1) | 2.59 (2.41x10^-1) |
| Pars Opercularis | 2.8 (1.46x10^-1) | 2.77 (1.49x10^-1) | 2.68 (1.13x10^-1) | 2.59 (2.21x10^-1) |
| Pars Orbitalis | 2.84 (1.95x10^-1) | 2.83 (2.64x10^-1) | 2.81 (1.37x10^-1) | 2.68 (3.19x10^-1) |
| Pars Triangularis | 2.63 (2.12x10^-1) | 2.62 (1.58x10^-1) | 2.61 (1.83x10^-1) | 2.36 (2.35x10^-1) |
| Pericalcarine | 1.73 (1.55x10^-1) | 1.75 (1.61x10^-1) | 1.73 (9.06x10^-2) | 1.64 (1.63x10^-1) |
| Postcentral | 2.27 (1.53x10^-1) | 2.22 (1.73x10^-1) | 2.26 (1.57x10^-1) | 2.23 (3.08x10^-1) |
| Posterior Cingulate | 2.54 (1.32x10^-1) | 2.5 (1.79x10^-1) | 2.52 (1.62x10^-1) | 2.42 (2.08x10^-1) |
| Precentral | 2.72 (1.28x10^-1) | 2.65 (1.77x10^-1) | 2.71 (1.46x10^-1) | 2.56 (1.68x10^-1) |
| Precuneus | 2.62 (1.02x10^-1) | 2.63 (1.51x10^-1) | 2.58 (1.05x10^-1) | 2.51 (1.53x10^-1) |
| Rostral Anterior Cingulate | 2.89 (1.86x10^-1) | 2.79 (2.27x10^-1) | 2.87 (2.64x10^-1) | 2.82 (1.28x10^-1) |
| Rostral Middle Frontal | 2.53 (1.38x10^-1) | 2.42 (1.35x10^-1) | 2.51 (1.35x10^-1) | 2.3 (1.95x10^-1) |

| | | | | |
|---|---|---|---|---|
| Superior Frontal | 2.85 (1.70x10^-1) | 2.78 (1.57x10^-1) | 2.83 (1.41x10^-1) | 2.61 (2.19x10^-1) |
| Superior Parietal | 2.41 (1.50x10^-1) | 2.4 (1.37x10^-1) | 2.38 (1.46x10^-1) | 2.33 (1.57x10^-1) |
| Superior Temporal | 3 (1.06x10^-1) | 3 (1.60x10^-1) | 2.96 (1.24x10^-1) | 2.92 (1.65x10^-1) |
| Supramarginal | 2.75 (1.14x10^-1) | 2.77 (1.94x10^-1) | 2.73 (1.38x10^-1) | 2.62 (1.67x10^-1) |
| Temporal Pole | 3.42 (2.00x10^-1) | 3.51 (4.05x10^-1) | 3.37 (2.97x10^-1) | 3.5 (5.69x10^-1) |
| Transverse Temporal | 2.66 (2.07x10^-1) | 2.7 (1.95x10^-1) | 2.53 (1.49x10^-1) | 2.5 (3.54x10^-1) |

LH = left hemisphere
RH = right hemisphere

**Table S4 General linear model results: Cortical thickness**

| **Region** | **Comparison** | ***p*-value** | ***Bayes factor*** |
|---|---|---|---|
| Banks of superior temporal sulcus | LC vs. LP | .99 | 8.33 |
| | RC vs. RP | .15 | 2.16 |
| | LP vs. RP | .40 | 2.81 |
| Caudal anterior cingulate | LC vs. LP | .99 | 8.32 |
| | RC vs. RP | .96 | 7.99 |
| | LP vs. RP | .29 | 1.65 |
| Caudal middle frontal | LC vs. LP | .99 | 8.30 |
| | RC vs. RP | **.02** | 4.61x10^-2 |
| | LP vs. RP | **.01** | 1.01x10^-2 |
| Cuneus | LC vs. LP | .99 | 8.34 |
| | RC vs. RP | **.02** | 1.05x10^-1 |
| | LP vs. RP | .29 | 2.24 |
| Entorhinal | LC vs. LP | .99 | 4.75 |
| | RC vs. RP | .82 | 7.63 |
| | LP vs. RP | .40 | 2.84 |
| Frontal pole | LC vs. LP | .99 | 6.88 |
| | RC vs. RP | **.02** | 2.35x10^-2 |
| | LP vs. RP | .18 | 5.53x10^-1 |
| Fusiform | LC vs. LP | .99 | 9.48x10^-1 |
| | RC vs. RP | **.02** | 4.92x10^-2 |
| | LP vs. RP | .55 | 4.45 |
| Inferior parietal | LC vs. LP | .99 | 8.15 |
| | RC vs. RP | .15 | 1.86 |
| | LP vs. RP | .76 | 5.36 |
| Inferior temporal | LC vs. LP | .99 | 2.83 |
| | RC vs. RP | .07 | 6.62x10^-1 |
| | LP vs. RP | .91 | 5.61 |
| Insula | LC vs. LP | .99 | 6.78 |
| | RC vs. RP | **.03** | 3.43x10^-1 |
| | LP vs. RP | .40 | 3.24 |
| Isthmus | LC vs. LP | .99 | 7.24 |
| | RC vs. RP | **.02** | 8.85x10^-2 |
| | LP vs. RP | .22 | 8.98x10^-1 |
| Lateral occipital | LC vs. LP | .99 | 7.11 |
| | RC vs. RP | .49 | 5.31 |
| | LP vs. RP | .52 | 4.00 |
| Lateral orbitofrontal | LC vs. LP | .99 | 7.73 |
| | RC vs. RP | **.02** | 1.36x10^-1 |
| | LP vs. RP | .22 | 1.02 |
| Lingual | LC vs. LP | .99 | 3.23 |
| | RC vs. RP | **.02** | 5.65x10^-2 |
| | LP vs. RP | .34 | 2.20 |

| | | | |
|---|---|---|---|
| Medial orbitofrontal | LC vs. LP | .99 | 8.04 |
| | RC vs. RP | .13 | 1.65 |
| | LP vs. RP | .18 | 5.00x10^-1 |
| Middle temporal | LC vs. LP | .99 | 8.07 |
| | RC vs. RP | **.03** | 3.14x10^-1 |
| | LP vs. RP | .40 | 3.27 |
| Parahippocampal | LC vs. LP | .99 | 4.50 |
| | RC vs. RP | .06 | 5.36x10^-1 |
| | LP vs. RP | .29 | 1.69 |
| Paracentral | LC vs. LP | .99 | 4.53 |
| | RC vs. RP | **.03** | 3.43x10^-1 |
| | LP vs. RP | .26 | 1.38 |
| Pars opercularis | LC vs. LP | .99 | 1.39 |
| | RC vs. RP | .06 | 5.12x10^-1 |
| | LP vs. RP | .19 | 6.85x10^-1 |
| Pars orbitalis | LC vs. LP | .99 | 5.07 |
| | RC vs. RP | .95 | 7.95 |
| | LP vs. RP | .55 | 4.27 |
| Pars triangularis | LC vs. LP | .99 | 8.07 |
| | RC vs. RP | .07 | 8.04x10^-1 |
| | LP vs. RP | .08 | 1.10x10^-1 |
| Pericalcarine | LC vs. LP | .99 | 8.08 |
| | RC vs. RP | .10 | 1.35 |
| | LP vs. RP | .18 | 2.66x10^-1 |
| Postcentral | LC vs. LP | .99 | 7.69 |
| | RC vs. RP | .90 | 7.84 |
| | LP vs. RP | .40 | 3.01 |
| Posterior cingulate | LC vs. LP | >.99 | 8.37 |
| | RC vs. RP | **.02** | 2.00x10^-1 |
| | LP vs. RP | .18 | 5.27x10^-1 |
| Precentral | LC vs. LP | .99 | 5.16 |
| | RC vs. RP | .28 | 3.19 |
| | LP vs. RP | .22 | 1.04 |
| Precuneus | LC vs. LP | .99 | 5.60 |
| | RC vs. RP | **.02** | 1.61x10^-1 |
| | LP vs. RP | .18 | 5.69x10^-1 |
| Rostral anterior cingulate | LC vs. LP | .99 | 5.12 |
| | RC vs. RP | .69 | 6.96 |
| | LP vs. RP | .29 | 1.81 |
| Rostral middle frontal | LC vs. LP | .99 | 8.20 |
| | RC vs. RP | **.02** | 3.75x10^-2 |
| | LP vs. RP | **<.001** | 1.06x10^-3 |
| Superior frontal | LC vs. LP | .99 | 8.13 |
| | RC vs. RP | **.03** | 1.84x10^-1 |
| | LP vs. RP | **<.001** | 2.65x10^-3 |
| Superior parietal | LC vs. LP | .99 | 8.15 |
| | RC vs. RP | .14 | 1.94 |
| | LP vs. RP | .29 | 1.88 |
| Superior temporal | LC vs. LP | .99 | 8.05 |
| | RC vs. RP | .28 | 3.24 |
| | LP vs. RP | .50 | 3.94 |
| Supramarginal | LC vs. LP | .99 | 5.66 |
| | RC vs. RP | .07 | 6.96x10^-1 |

| | | | |
|---|---|---|---|
| | LP vs. RP | .22 | 8.24x10^-1 |
| Temporal pole | LC vs. LP | >.99 | 8.37 |
| | RC vs. RP | .69 | 6.90 |
| | LP vs. RP | .40 | 3.09 |
| Transverse temporal | LC vs. LP | .99 | 7.45 |
| | RC vs. RP | **.02** | 1.50x10^-2 |
| | LP vs. RP | .22 | 1.08 |

LC = left hemisphere of controls
LP = left hemisphere of patients
RC = right hemisphere of controls
RP = right hemisphere of patients
*p*-values less than .05 are bolded.

**Table S5 Descriptive statistics: Median normalized cortical surface area (median absolute deviations per cortical area**

| Region | Control LH | Control RH | Patient LH | Patient RH |
|---|---|---|---|---|
| Banks of Superior Temporal Sulcus | 4.13x10^-1 (5.34x10^-2) | 3.65x10^-1 (3.72x10^-2) | 4.08x10^-1 (5.47x10^-2) | 3.48x10^-1 (5.92x10^-2) |
| Caudal Anterior Cingulate | 2.35x10^-1 (4.14x10^-2) | 2.99x10^-1 (4.80x10^-2) | 2.39x10^-1 (4.03x10^-2) | 2.54x10^-1 (7.73x10^-2) |
| Caudal Middle Frontal | 8.51x10^-1 (7.57x10^-2) | 7.82x10^-1 (1.30x10^-1) | 9.04x10^-1 (1.24x10^-1) | 7.61x10^-1 (1.27x10^-1) |
| Cuneus | 6.34x10^-1 (8.12x10^-2) | 6.56x10^-1 (6.47x10^-2) | 6.13x10^-1 (6.46x10^-2) | 6.75x10^-1 (5.51x10^-2) |
| Entorhinal | 1.70x10^-1 (2.21x10^-2) | 1.59x10^-1 (3.39x10^-2) | 1.52x10^-1 (3.27x10^-2) | 1.62x10^-1 (1.98x10^-2) |
| Frontal Pole | 1.14x10^-1 (1.05x10^-2) | 1.42x10^-1 (1.51x10^-2) | 1.13x10^-1 (9.08x10^-3) | 1.43x10^-1 (2.01x10^-2) |
| Fusiform | 1.23 (9.00x10^-2) | 1.24 (1.09x10^-1) | 1.26 (1.53x10^-1) | 1.31 (8.90x10^-2) |
| Inferior Parietal | 1.82 (1.86x10^-1) | 2.14 (2.24x10^-1) | 1.79 (1.42x10^-1) | 2.2 (2.73x10^-1) |
| Inferior Temporal | 1.39 (1.48x10^-1) | 1.31 (1.19x10^-1) | 1.4 (1.93x10^-1) | 1.37 (9.34x10^-2) |
| Insula | 8.57x10^-1 (5.92x10^-2) | 8.27x10^-1 (6.29x10^-2) | 8.97x10^-1 (9.83x10^-2) | 8.73x10^-1 (6.95x10^-2) |
| Isthmus | 4.10x10^-1 (5.17x10^-2) | 3.67x10^-1 (3.49x10^-2) | 4.12x10^-1 (6.16x10^-2) | 3.76x10^-1 (4.60x10^-2) |
| Lateral Occipital | 2.02 (2.10x10^-1) | 2.1 (1.76x10^-1) | 2.01 (1.80x10^-1) | 2.05 (2.31x10^-1) |
| Lateral Orbitofrontal | 1.07 (6.76x10^-2) | 1.02 (8.41x10^-2) | 1.07 (9.37x10^-2) | 1.09 (7.28x10^-2) |
| Lingual | 1.25 (1.36x10^-1) | 1.34 (1.52x10^-1) | 1.26 (1.18x10^-1) | 1.32 (1.57x10^-1) |
| Medial Orbitofrontal | 7.58x10^-1 (5.75x10^-2) | 8.13x10^-1 (4.96x10^-2) | 7.75x10^-1 (5.79x10^-2) | 8.52x10^-1 (4.46x10^-2) |
| Middle Temporal | 1.33 (8.11x10^-2) | 1.45 (8.40x10^-2) | 1.35 (7.67x10^-2) | 1.41 (4.05x10^-2) |
| Parahippocampal | 5.01x10^-1 (4.98x10^-2) | 5.65x10^-1 (4.66x10^-2) | 5.09x10^-1 (6.97x10^-2) | 5.99x10^-1 (1.05x10^-1) |
| Paracentral | 2.49x10^-1 (2.69x10^-2) | 2.37x10^-1 (2.53x10^-2) | 2.42x10^-1 (3.98x10^-2) | 2.59x10^-1 (2.92x10^-2) |
| Pars Opercularis | 6.51x10^-1 (8.54x10^-2) | 5.27x10^-1 (5.57x10^-2) | 6.52x10^-1 (5.83x10^-2) | 5.06x10^-1 (4.28x10^-2) |
| Pars Orbitalis | 2.88x10^-1 (1.80x10^-2) | 3.43x10^-1 (3.24x10^-2) | 2.82x10^-1 (2.71x10^-2) | 3.55x10^-1 (2.40x10^-2) |
| Pars Triangularis | 5.60x10^-1 (6.30x10^-2) | 6.19x10^-1 (9.88x10^-2) | 5.40x10^-1 (7.78x10^-2) | 6.07x10^-1 (8.17x10^-2) |
| Pericalcarine | 5.73x10^-1 (8.84x10^-2) | 6.31x10^-1 (9.16x10^-2) | 5.77x10^-1 (9.78x10^-2) | 6.56x10^-1 (8.94x10^-2) |
| Postcentral | 1.57 (1.38x10^-1) | 1.5 (1.62x10^-1) | 1.56 (1.12x10^-1) | 1.55 (1.88x10^-1) |
| Posterior Cingulate | 4.64x10^-1 (4.12x10^-2) | 4.76x10^-1 (5.24x10^-2) | 4.95x10^-1 (5.13x10^-2) | 4.78x10^-1 (6.62x10^-2) |
| Precentral | 1.81 (1.18x10^-1) | 1.81 (2.23x10^-1) | 1.82 (1.57x10^-1) | 1.8 (1.28x10^-1) |
| Precuneus | 1.5 (1.06x10^-1) | 1.57 (1.40x10^-1) | 1.56 (1.32x10^-1) | 1.52 (1.08x10^-1) |
| Rostral Anterior Cingulate | 3.23x10^-1 (6.06x10^-2) | 2.47x10^-1 (3.67x10^-2) | 3.24x10^-1 (4.39x10^-2) | 2.42x10^-1 (8.91x10^-2) |
| Rostral Middle Frontal | 2.33 (1.90x10^-1) | 2.41 (1.52x10^-1) | 2.27 (1.65x10^-1) | 2.23 (2.86x10^-1) |
| Superior Frontal | 2.83 (1.49x10^-1) | 2.72 (2.39x10^-1) | 2.91 (2.96x10^-1) | 2.65 (2.26x10^-1) |

| | | | | |
|---|---|---|---|---|
| Superior Parietal | 2.13 (2.33x10^-1) | 2.05 (2.65x10^-1) | 2.23 (2.64x10^-1) | 2.11 (1.45x10^-1) |
| Superior Temporal | 1.58 (9.30x10^-2) | 1.45 (1.02x10^-1) | 1.53 (1.12x10^-1) | 1.42 (1.35x10^-1) |
| Supramarginal | 1.54 (1.96x10^-1) | 1.42 (1.72x10^-1) | 1.53 (2.29x10^-1) | 1.57 (1.24x10^-1) |
| Temporal Pole | 2.01x10^-1 (2.81x10^-2) | 1.93x10^-1 (1.84x10^-2) | 1.97x10^-1 (1.66x10^-2) | 2.06x10^-1 (2.80x10^-2) |
| Transverse Temporal | 1.76x10^-1 (1.55x10^-2) | 1.30x10^-1 (1.17x10^-2) | 1.71x10^-1 (2.63x10^-2) | 1.27x10^-1 (2.51x10^-2) |

LH = left hemisphere
RH = right hemisphere

**Table S6 General linear model results: Cortical surface area**

| Region | Comparison | *p*-value | *Bayes factor* |
|---|---|---|---|
| Banks of superior temporal sulcus | LC vs. LP | .54 | 0.98 |
| | RC vs. RP | .87 | 5.79 |
| | LP vs. RP | .10 | 3.59x10^-1 |
| Caudal anterior cingulate | LC vs. LP | .95 | 7.93 |
| | RC vs. RP | .29 | 1.26 |
| | LP vs. RP | .77 | 4.68 |
| Caudal middle frontal | LC vs. LP | .95 | 7.31 |
| | RC vs. RP | .91 | 7.86 |
| | LP vs. RP | .31 | 1.45 |
| Cuneus | LC vs. LP | .95 | 8.28 |
| | RC vs. RP | .87 | 6.16 |
| | LP vs. RP | .39 | 2.28 |
| Entorhinal | LC vs. LP | .54 | 1.82 |
| | RC vs. RP | .92 | 7.96 |
| | LP vs. RP | .97 | 5.65 |
| Frontal pole | LC vs. LP | .54 | 1.61 |
| | RC vs. RP | .87 | 7.44 |
| | LP vs. RP | **<.001** | 4.82x10^-6 |
| Fusiform | LC vs. LP | .95 | 7.72 |
| | RC vs. RP | .87 | 7.05 |
| | LP vs. RP | .87 | 5.40 |
| Inferior parietal | LC vs. LP | .81 | 5.73 |
| | RC vs. RP | .91 | 7.70 |
| | LP vs. RP | **<.001** | 4.62x10^-3 |
| Inferior temporal | LC vs. LP | .86 | 6.62 |
| | RC vs. RP | .87 | 7.39 |
| | LP vs. RP | .39 | 2.30 |
| Insula | LC vs. LP | .86 | 6.49 |
| | RC vs. RP | .87 | 5.74 |
| | LP vs. RP | .77 | 4.53 |
| Isthmus | LC vs. LP | .81 | 4.85 |
| | RC vs. RP | .87 | 7.31 |
| | LP vs. RP | .09 | 2.61x10^-1 |
| Lateral occipital | LC vs. LP | .97 | 8.36 |
| | RC vs. RP | .87 | 6.35 |
| | LP vs. RP | .87 | 5.43 |
| Lateral orbitofrontal | LC vs. LP | .95 | 8.23 |
| | RC vs. RP | **.02** | 4.88x10^-2 |
| | LP vs. RP | .49 | 2.92 |
| Lingual | LC vs. LP | .95 | 8.21 |
| | RC vs. RP | .91 | 7.84 |
| | LP vs. RP | .61 | 3.54 |

| | | | |
|---|---|---|---|
| Medial orbitofrontal | LC vs. LP | .58 | 2.35 |
| | RC vs. RP | .87 | 6.54 |
| | LP vs. RP | .19 | 6.95x10^-1 |
| Middle temporal | LC vs. LP | .81 | 5.63 |
| | RC vs. RP | .73 | 4.06 |
| | LP vs. RP | .17 | 6.31x10^-1 |
| Parahippocampal | LC vs. LP | .95 | 7.54 |
| | RC vs. RP | **.02** | 4.39x10^-2 |
| | LP vs. RP | **<.001** | 5.38x10^-3 |
| Paracentral | LC vs. LP | .95 | 8.30 |
| | RC vs. RP | **<.001** | 5.88x10^-2 |
| | LP vs. RP | .39 | 2.30 |
| Pars opercularis | LC vs. LP | .81 | 6.17 |
| | RC vs. RP | .59 | 2.87 |
| | LP vs. RP | **<.001** | 2.36x10^-5 |
| Pars orbitalis | LC vs. LP | .58 | 2.60 |
| | RC vs. RP | .29 | 1.07 |
| | LP vs. RP | **<.001** | 5.96x10^-8 |
| Pars triangularis | LC vs. LP | .95 | 8.18 |
| | RC vs. RP | .70 | 3.79 |
| | LP vs. RP | .39 | 2.22 |
| Pericalcarine | LC vs. LP | .95 | 8.12 |
| | RC vs. RP | .92 | 7.92 |
| | LP vs. RP | .39 | 2.01 |
| Postcentral | LC vs. LP | .81 | 5.36 |
| | RC vs. RP | .87 | 5.62 |
| | LP vs. RP | .87 | 5.34 |
| Posterior cingulate | LC vs. LP | .81 | 5.35 |
| | RC vs. RP | .87 | 7.43 |
| | LP vs. RP | .87 | 5.47 |
| Precentral | LC vs. LP | .81 | 3.76 |
| | RC vs. RP | .87 | 5.74 |
| | LP vs. RP | .95 | 5.61 |
| Precuneus | LC vs. LP | .92 | 7.08 |
| | RC vs. RP | .87 | 5.54 |
| | LP vs. RP | .87 | 5.22 |
| Rostral anterior cingulate | LC vs. LP | .81 | 5.86 |
| | RC vs. RP | .87 | 7.58 |
| | LP vs. RP | **<.001** | 5.14x10^-3 |
| Rostral middle frontal | LC vs. LP | .95 | 8.06 |
| | RC vs. RP | .14 | 3.36 |
| | LP vs. RP | .87 | 5.10 |
| Superior frontal | LC vs. LP | .81 | 3.89 |
| | RC vs. RP | .87 | 7.51 |
| | LP vs. RP | .12 | 4.17x10^-1 |
| Superior parietal | LC vs. LP | .81 | 4.69 |
| | RC vs. RP | .91 | 7.82 |
| | LP vs. RP | .69 | 3.83 |
| Superior temporal | LC vs. LP | .20 | 8.12x10^-2 |
| | RC vs. RP | .14 | 5.88x10^-1 |
| | LP vs. RP | .07 | 1.51x10^-1 |
| Supramarginal | LC vs. LP | .54 | 1.78 |
| | RC vs. RP | .14 | 4.18x10^-1 |
| | LP vs. RP | .87 | 5.28 |

| | | | |
|---|---|---|---|
| Temporal pole | LC vs. LP | .97 | 8.34 |
| | RC vs. RP | .59 | 2.95 |
| | LP vs. RP | .87 | 5.47 |
| Transverse temporal | LC vs. LP | .20 | 4.64x10^-1 |
| | RC vs. RP | .87 | 6.76 |
| | LP vs. RP | **<.001** | 1.42x10^-5 |

LC = left hemisphere of controls
LP = left hemisphere of patients
RC = right hemisphere of controls
RP = right hemisphere of patients
*p*-values less than .05 are bolded.

**Table S7 Descriptive statistics: Median normalized region volume (median absolute deviations) per cortical area**

| Region | Control LH | Control RH | Patient LH | Patient RH |
|---|---|---|---|---|
| Banks of Superior Temporal Sulcus | 5.96x10^-3 (1.10x10^-3) | 5.25x10^-3 (7.56x10^-4) | 5.58x10^-3 (9.94x10^-4) | 4.74x10^-3 (6.36x10^-4) |
| Caudal Anterior Cingulate | 3.71x10^-3 (6.67x10^-4) | 4.62x10^-3 (1.01x10^-3) | 3.82x10^-3 (9.24x10^-4) | 3.89x10^-3 (1.58x10^-3) |
| Caudal Middle Frontal | 1.37x10^-2 (1.50x10^-3) | 1.25x10^-2 (2.38x10^-3) | 1.45x10^-2 (2.77x10^-3) | 1.11x10^-2 (1.37x10^-3) |
| Cuneus | 7.40x10^-3 (1.16x10^-3) | 8.04x10^-3 (1.23x10^-3) | 7.25x10^-3 (2.17x10^-3) | 8.40x10^-3 (6.12x10^-4) |
| Entorhinal | 3.86x10^-3 (5.96x10^-4) | 3.71x10^-3 (8.79x10^-4) | 3.58x10^-3 (4.69x10^-4) | 3.73x10^-3 (1.36x10^-3) |
| Frontal Pole | 2.63x10^-3 (4.52x10^-4) | 3.27x10^-3 (4.02x10^-4) | 2.44x10^-3 (5.64x10^-4) | 2.86x10^-3 (5.00x10^-4) |
| Fusiform | 2.16x10^-2 (1.62x10^-3) | 2.29x10^-2 (2.45x10^-3) | 2.17x10^-2 (2.95x10^-3) | 2.22x10^-2 (2.59x10^-3) |
| Inferior Parietal | 2.96x10^-2 (4.09x10^-3) | 3.69x10^-2 (5.81x10^-3) | 2.88x10^-2 (3.15x10^-3) | 3.75x10^-2 (3.96x10^-3) |
| Inferior Temporal | 2.65x10^-2 (2.97x10^-3) | 2.59x10^-2 (3.32x10^-3) | 2.62x10^-2 (3.59x10^-3) | 2.53x10^-2 (3.01x10^-3) |
| Insula | 1.48x10^-2 (1.03x10^-3) | 1.46x10^-2 (1.44x10^-3) | 1.50x10^-2 (1.47x10^-3) | 1.49x10^-2 (1.52x10^-3) |
| Isthmus | 6.24x10^-3 (6.96x10^-4) | 5.76x10^-3 (7.47x10^-4) | 6.25x10^-3 (1.41x10^-3) | 5.23x10^-3 (6.95x10^-4) |
| Lateral Occipital | 2.87x10^-2 (3.19x10^-3) | 3.00x10^-2 (2.83x10^-3) | 2.87x10^-2 (2.67x10^-3) | 3.05x10^-2 (2.36x10^-3) |
| Lateral Orbitofrontal | 1.79x10^-2 (1.37x10^-3) | 1.71x10^-2 (1.23x10^-3) | 1.77x10^-2 (1.15x10^-3) | 1.71x10^-2 (9.44x10^-4) |
| Lingual | 1.56x10^-2 (2.00x10^-3) | 1.68x10^-2 (2.51x10^-3) | 1.51x10^-2 (2.95x10^-3) | 1.65x10^-2 (2.09x10^-3) |
| Medial Orbitofrontal | 1.18x10^-2 (1.15x10^-3) | 1.35x10^-2 (1.18x10^-3) | 1.26x10^-2 (8.73x10^-4) | 1.30x10^-2 (3.87x10^-4) |
| Middle Temporal | 2.80x10^-2 (3.68x10^-3) | 3.03x10^-2 (2.20x10^-3) | 2.85x10^-2 (2.30x10^-3) | 2.79x10^-2 (1.24x10^-3) |
| Parahippocampal | 8.17x10^-3 (1.07x10^-3) | 8.87x10^-3 (7.65x10^-4) | 8.09x10^-3 (1.08x10^-3) | 8.78x10^-3 (7.56x10^-4) |
| Paracentral | 4.42x10^-3 (6.49x10^-4) | 4.22x10^-3 (5.31x10^-4) | 4.31x10^-3 (7.23x10^-4) | 4.25x10^-3 (2.40x10^-4) |
| Pars Opercularis | 1.12x10^-2 (1.70x10^-3) | 8.97x10^-3 (1.05x10^-3) | 1.13x10^-2 (1.43x10^-3) | 8.28x10^-3 (1.41x10^-3) |
| Pars Orbitalis | 6.00x10^-3 (5.57x10^-4) | 6.95x10^-3 (7.75x10^-4) | 5.70x10^-3 (1.08x10^-3) | 7.09x10^-3 (1.20x10^-3) |
| Pars Triangularis | 9.05x10^-3 (1.24x10^-3) | 1.03x10^-2 (2.11x10^-3) | 8.96x10^-3 (1.28x10^-3) | 9.61x10^-3 (2.15x10^-3) |
| Pericalcarine | 4.67x10^-3 (1.03x10^-3) | 5.30x10^-3 (1.17x10^-3) | 4.88x10^-3 (1.05x10^-3) | 4.92x10^-3 (4.81x10^-4) |
| Postcentral | 2.19x10^-2 (2.52x10^-3) | 2.09x10^-2 (1.88x10^-3) | 2.27x10^-2 (3.05x10^-3) | 2.09x10^-2 (3.52x10^-3) |
| Posterior Cingulate | 7.02x10^-3 (9.60x10^-4) | 7.39x10^-3 (7.71x10^-4) | 7.04x10^-3 (7.81x10^-4) | 6.91x10^-3 (1.18x10^-3) |
| Precentral | 2.93x10^-2 (2.32x10^-3) | 2.84x10^-2 (2.95x10^-3) | 3.09x10^-2 (3.63x10^-3) | 2.79x10^-2 (3.95x10^-3) |
| Precuneus | 2.31x10^-2 (2.53x10^-3) | 2.40x10^-2 (2.50x10^-3) | 2.38x10^-2 (2.27x10^-3) | 2.24x10^-2 (2.60x10^-3) |
| Rostral Anterior Cingulate | 5.94x10^-3 (8.95x10^-4) | 4.45x10^-3 (9.42x10^-4) | 5.93x10^-3 (8.32x10^-4) | 4.79x10^-3 (3.52x10^-4) |
| Rostral Middle Frontal | 3.70x10^-2 (3.57x10^-3) | 3.82x10^-2 (4.22x10^-3) | 3.57x10^-2 (3.66x10^-3) | 3.30x10^-2 (6.44x10^-3) |

| | | | | |
|---|---|---|---|---|
| Superior Frontal | 5.22x10^-2 (4.79x10^-3) | 4.86x10^-2 (5.25x10^-3) | 5.26x10^-2 (7.14x10^-3) | 4.42x10^-2 (3.19x10^-3) |
| Superior Parietal | 3.08x10^-2 (5.14x10^-3) | 3.04x10^-2 (3.92x10^-3) | 3.24x10^-2 (4.33x10^-3) | 3.02x10^-2 (2.96x10^-3) |
| Superior Temporal | 3.00x10^-2 (2.53x10^-3) | 2.77x10^-2 (2.78x10^-3) | 2.94x10^-2 (2.56x10^-3) | 2.65x10^-2 (3.12x10^-3) |
| Supramarginal | 2.56x10^-2 (3.74x10^-3) | 2.36x10^-2 (3.49x10^-3) | 2.57x10^-2 (4.49x10^-3) | 2.56x10^-2 (4.01x10^-3) |
| Temporal Pole | 5.23x10^-3 (7.17x10^-4) | 5.21x10^-3 (8.66x10^-4) | 5.41x10^-3 (1.06x10^-3) | 5.88x10^-3 (6.90x10^-4) |
| Transverse Temporal | 2.80x10^-3 (4.28x10^-4) | 2.17x10^-3 (2.24x10^-4) | 2.61x10^-3 (2.42x10^-4) | 1.93x10^-3 (3.26x10^-4) |

LH = left hemisphere
RH = right hemisphere

**Table S8 General linear model results: Cortical volume**

| **Region** | **Comparison** | ***p*-value** | ***Bayes factor*** |
|---|---|---|---|
| Banks of superior temporal sulcus | LC vs. LP | .75 | 2.61 |
| | RC vs. RP | .61 | 5.78 |
| | LP vs. RP | .27 | 1.30 |
| Caudal anterior cingulate | LC vs. LP | .99 | 8.36 |
| | RC vs. RP | .30 | 1.64 |
| | LP vs. RP | .84 | 4.97 |
| Caudal middle frontal | LC vs. LP | .75 | 5.36 |
| | RC vs. RP | .61 | 5.66 |
| | LP vs. RP | .07 | 1.66x10^-1 |
| Cuneus | LC vs. LP | .75 | 5.67 |
| | RC vs. RP | .84 | 7.57 |
| | LP vs. RP | .84 | 4.61 |
| Entorhinal | LC vs. LP | .75 | 1.16 |
| | RC vs. RP | .85 | 7.67 |
| | LP vs. RP | .45 | 2.12 |
| Frontal pole | LC vs. LP | .75 | 2.81 |
| | RC vs. RP | .30 | 5.04x10^-1 |
| | LP vs. RP | .07 | 1.92x10^-1 |
| Fusiform | LC vs. LP | .99 | 8.33 |
| | RC vs. RP | .53 | 4.70 |
| | LP vs. RP | .97 | 5.63 |
| Inferior parietal | LC vs. LP | .75 | 5.72 |
| | RC vs. RP | .79 | 7.07 |
| | LP vs. RP | **.01** | 2.29x10^-3 |
| Inferior temporal | LC vs. LP | .99 | 7.77 |
| | RC vs. RP | .79 | 7.24 |
| | LP vs. RP | .80 | 3.54 |
| Insula | LC vs. LP | .99 | 7.51 |
| | RC vs. RP | .83 | 7.45 |
| | LP vs. RP | .97 | 5.65 |
| Isthmus | LC vs. LP | .75 | 5.40 |
| | RC vs. RP | .30 | 7.72x10^-1 |
| | LP vs. RP | .07 | 1.83x10^-1 |
| Lateral occipital | LC vs. LP | .99 | 8.36 |
| | RC vs. RP | .73 | 6.53 |
| | LP vs. RP | .84 | 4.72 |
| Lateral orbitofrontal | LC vs. LP | .99 | 7.39 |
| | RC vs. RP | .30 | 1.27 |
| | LP vs. RP | .83 | 4.07 |
| Lingual | LC vs. LP | .77 | 6.25 |

| | | | |
|---|---|---|---|
| | RC vs. RP | .45 | 3.81 |
| | LP vs. RP | .93 | 5.37 |
| Medial orbitofrontal | LC vs. LP | .75 | 4.39 |
| | RC vs. RP | .61 | 5.56 |
| | LP vs. RP | .12 | 6.06x10^-1 |
| Middle temporal | LC vs. LP | .75 | 3.14 |
| | RC vs. RP | .30 | 1.36 |
| | LP vs. RP | .97 | 5.64 |
| Parahippocampal | LC vs. LP | .99 | 8.36 |
| | RC vs. RP | .34 | 2.77 |
| | LP vs. RP | .07 | 2.39x10^-1 |
| Paracentral | LC vs. LP | .99 | 8.34 |
| | RC vs. RP | .34 | 2.85 |
| | LP vs. RP | .93 | 5.42 |
| Pars opercularis | LC vs. LP | .75 | 5.70 |
| | RC vs. RP | .30 | 2.16 |
| | LP vs. RP | **<.001** | 3.55x10^-5 |
| Pars orbitalis | LC vs. LP | .75 | 3.40 |
| | RC vs. RP | .30 | 2.43 |
| | LP vs. RP | **<.001** | 2.34x10^-4 |
| Pars triangularis | LC vs. LP | .99 | 8.36 |
| | RC vs. RP | .30 | 2.05 |
| | LP vs. RP | .93 | 5.51 |
| Pericalcarine | LC vs. LP | .99 | 8.36 |
| | RC vs. RP | .54 | 5.04 |
| | LP vs. RP | .97 | 5.56 |
| Postcentral | LC vs. LP | .99 | 7.80 |
| | RC vs. RP | .52 | 4.49 |
| | LP vs. RP | .84 | 4.99 |
| Posterior cingulate | LC vs. LP | .99 | 7.80 |
| | RC vs. RP | .30 | 1.49 |
| | LP vs. RP | .81 | 3.80 |
| Precentral | LC vs. LP | .75 | 2.11 |
| | RC vs. RP | .98 | 7.99 |
| | LP vs. RP | .26 | 1.04 |
| Precuneus | LC vs. LP | .77 | 6.09 |
| | RC vs. RP | .30 | 1.56 |
| | LP vs. RP | .84 | 4.48 |
| Rostral anterior cingulate | LC vs. LP | .99 | 8.37 |
| | RC vs. RP | .98 | 8.00 |
| | LP vs. RP | **<.001** | 5.80x10^-3 |
| Rostral middle frontal | LC vs. LP | .99 | 8.33 |
| | RC vs. RP | **<.001** | 1.49x10^-3 |
| | LP vs. RP | .09 | 3.07x10^-1 |
| Superior frontal | LC vs. LP | .75 | 3.52 |
| | RC vs. RP | .30 | 2.08 |
| | LP vs. RP | **.01** | 1.39x10^-2 |
| Superior parietal | LC vs. LP | .75 | 4.66 |
| | RC vs. RP | .88 | 7.81 |
| | LP vs. RP | .84 | 4.20 |
| Superior temporal | LC vs. LP | .75 | 2.56 |
| | RC vs. RP | .30 | 1.76 |
| | LP vs. RP | .07 | 2.31x10^-1 |
| Supramarginal | LC vs. LP | .75 | 5.50 |

| | | | |
|---|---|---|---|
| | RC vs. RP | .30 | 2.29 |
| | LP vs. RP | .84 | 4.91 |
| Temporal pole | LC vs. LP | .75 | 5.20 |
| | RC vs. RP | .30 | 1.61 |
| | LP vs. RP | .87 | 5.13 |
| Transverse temporal | LC vs. LP | .75 | 2.39 |
| | RC vs. RP | .30 | 6.01x10^-1 |
| | LP vs. RP | **<.001** | 1.30x10^-4 |

LC = left hemisphere of controls
LP = left hemisphere of patients
RC = right hemisphere of controls
RP = right hemisphere of patients
*p*-values less than .05 are bolded.

**Table S9 Descriptive statistics: Median volume (median absolute deviations) per subcortical area**

| Region | Control LH | Control RH | Patient LH | Patient RH |
|---|---|---|---|---|
| Accumbens | 1.14x10^-3 (2.33x10^-4) | 1.22x10^-3 (1.97x10^-4) | 1.01x10^-3 (2.33x10^-4) | 1.14x10^-3 (1.65x10^-4) |
| Amygdala | 3.43x10^-3 (3.95x10^-4) | 3.60x10^-3 (3.29x10^-4) | 3.45x10^-3 (5.90x10^-4) | 3.79x10^-3 (4.80x10^-4) |
| Caudate | 7.89x10^-3 (7.50x10^-4) | 8.23x10^-3 (6.95x10^-4) | 7.56x10^-3 (6.59x10^-4) | 8.29x10^-3 (6.34x10^-4) |
| Cerebellum | 1.17x10^-1 (9.48x10^-3) | 1.19x10^-1 (8.94x10^-3) | 1.18x10^-1 (1.35x10^-2) | 1.23x10^-1 (1.19x10^-2) |
| Hippocampus | 8.20x10^-3 (5.78x10^-4) | 8.66x10^-3 (7.55x10^-4) | 8.22x10^-3 (9.93x10^-4) | 7.84x10^-3 (1.17x10^-3) |
| Pallidum | 4.13x10^-3 (2.32x10^-4) | 3.94x10^-3 (4.67x10^-4) | 3.78x10^-3 (6.03x10^-4) | 3.98x10^-3 (6.22x10^-4) |
| Putamen | 1.11x10^-2 (1.11x10^-3) | 1.14x10^-2 (9.41x10^-4) | 1.00x10^-2 (9.84x10^-4) | 1.13x10^-2 (1.40x10^-3) |
| Thalamus | 1.60x10^-2 (1.26x10^-3) | 1.59x10^-2 (9.39x10^-4) | 1.57x10^-2 (1.76x10^-3) | 1.60x10^-2 (1.81x10^-3) |
| Ventral Diencephalon | 7.77x10^-3 (6.66x10^-4) | 8.01x10^-3 (7.78x10^-4) | 8.01x10^-3 (9.78x10^-4) | 8.34x10^-3 (1.03x10^-3) |

LH = left hemisphere
RH = right hemisphere

**Table S10 General linear model results: Subcortical volume**

| Region | Comparison | *p*-value | *Bayes factor* |
|---|---|---|---|
| Accumbens | LC vs. LP | **.01** | 1.37x10^-2 |
| | RC vs. RP | **<.001** | 1.84x10^-2 |
| | LP vs. RP | .62 | 3.25 |
| Amygdala | LC vs. LP | .38 | 3.90 |
| | RC vs. RP | .52 | 2.99 |
| | LP vs. RP | .22 | 1.15 |
| Caudate | LC vs. LP | **.02** | 1.79x10^-1 |
| | RC vs. RP | .99 | 7.99 |
| | LP vs. RP | **<.01** | 3.28x10^-2 |
| Cerebellum | LC vs. LP | .56 | 6.31 |
| | RC vs. RP | .93 | 7.59 |
| | LP vs. RP | .66 | 4.34 |
| Hippocampus | LC vs. LP | .77 | 7.80 |
| | RC vs. RP | **.01** | 4.75x10^-2 |
| | LP vs. RP | .62 | 3.84 |
| Pallidum | LC vs. LP | **<.001** | 9.38x10^-3 |
| | RC vs. RP | .93 | 6.07 |
| | LP vs. RP | .22 | 1.06 |
| Putamen | LC vs. LP | **<.001** | 6.32x10^-4 |
| | RC vs. RP | .93 | 6.99 |
| | LP vs. RP | **<.01** | 1.49x10^-3 |

| | | | |
|---|---|---|---|
| Thalamus | LC vs. LP | .99 | 8.37 |
| | RC vs. RP | .93 | 7.37 |
| | LP vs. RP | .97 | 5.65 |
| Ventral diencephalon | LC vs. LP | .55 | 5.39 |
| | RC vs. RP | .99 | 8.00 |
| | LP vs. RP | .91 | 5.43 |

LC = left hemisphere of controls
LP = left hemisphere of patients
RC = right hemisphere of controls
RP = right hemisphere of patients
*p*-values less than .05 are bolded.

**Supplementary Material References**


1. Fox J, Weisberg S. *An {R} Companion to Applied Regression*. 3rd ed. Thousand Oaks, CA: Sage; 2019.

2. Signorell A. *DescTools: Tools for Descriptive Statistics*.; 2021.

3. Bates D, Mächler M, Bolker B, Walker S. Fitting linear mixed-effects models using lme4. *J Stat Softw*. 2015;67(1):1-48. doi:10.18637/jss.v067.i01

4. Fortin J-P. *NeuroCombat: Harmonization of Multi-Site Imaging Data with ComBat*.; 2021.

5. Revelle W. *Psych: Procedures for Personality and Psychological Research*. Evanston, Illinois, USA: Northwestern Unviersity; 2021.

6. Wickham H. *Stringr: Simple, Consistent Wrappers for Common String Operations*.; 2019.

7. Wickham H, Averick M, Bryan J, et al. Welcome to the tidyverse. *JOSS*. 2019;4(43):1686. doi:10.21105/joss.01686
